# Functional reorganization of sensory processing following long-term neural adaptation to optical defects

**DOI:** 10.1101/2020.02.19.956219

**Authors:** Antoine Barbot, Woon-Ju Park, Ru-Yuan Zhang, Krystel R Huxlin, Duje Tadin, Geunyoung Yoon

## Abstract

How we see is fundamentally limited by the eye’s optics, which determine retinal image quality and constrain neural processing. Elucidating how long-term exposure to optical defects alters visual processing is vital for understanding the human brain’s capacity for and limits of sensory plasticity. Using adaptive optics to bypass the eye’s optical aberrations, we assessed changes in visual processing in neurotypically-developed adults with keratoconus (KC)—a corneal disease causing severe optical aberrations during adulthood that cannot be fully corrected using conventional methods. As a result, KC patients are chronically exposed to degraded retinal images in their everyday life, making them an ideal model to understand how prolonged exposure to poor optical quality alters visual processing. Here, we show that when tested under similar fully-corrected optical conditions as neurotypical observers, KC patients exhibited altered contrast sensitivity, with impaired sensitivity for fine spatial details and better sensitivity for coarse spatial details. Both gains and losses in contrast sensitivity were more pronounced in patients with poorer habitual optical quality. Moreover, using an equivalent noise paradigm and a computational model of visual processing, we show that these alterations in visual processing are mediated by changes in signal enhancement of spatial frequency selective mechanisms. The present findings uncover fundamental properties of neural compensation mechanisms in response to long-term exposure to optical defects, which alter sensory processing and limit the benefits of improved optics. The outcome is a large-scale functional reorganization favoring the processing of sensory information less affected by the eye’s optics.

**Significance statement:** The eye’s optics represent an intrinsic limit to human visual perception, determining the quality of retinal images. Neural adaptation optimizes the brain’s limited sensory processing capacity to the structure of the degraded retinal inputs, providing an exceptional quality of vision given these optical limitations. Here, we show that prolonged exposure to poor optical quality results in a functional reorganization of visual processing that favors sensory information less affected by the eye’s optics. The present study helps elucidate how optical factors shape the way the brain processes visual information. Notably, the resulting adaptive neural plasticity limits the immediate perceptual benefits of optical interventions, a factor that must be taken into consideration when treating the increasing human population affected by optical defects.

---

Understanding how we see requires insights into the contribution of both optical and neural factors mediating visual perception, from the processing of images formed on the retina to the resulting perceptual representations. Visual processing is fundamentally limited by the eye’s optics, which determine retinal image quality and constrain performance. Human optics, however, are not fixed and can substantially change over our lifespan and in disease (1). Consequently, visual processing must be able to adapt to changes in sensory inputs over a range of different timescales and magnitudes. For instance, there is growing evidence that neural processing adapts to extended optical blur exposures, resulting in improved visual representations (2). Neural adaptation mechanisms can compensate for blur-induced reductions in physical contrast of high spatial frequency (SF) retinal signals (2-4). Even observers with typical optical quality show evidence of neural adaptation to their own optical blur, such that even modest changes in optical quality result in degraded subjective image quality (5).

Distinct neural adaptation mechanisms operate over different timescales (6,7). Longer exposure to a given environment induces longer-lasting effects that are mediated by separate mechanisms from the ones controlling short-term adaptation (6). Most studies investigated the effects of blur adaptation over relatively short-term periods (and often low magnitude of blur), thus limiting our understanding of how the adult human brain adapts to large changes in optical quality over long-term periods (i.e., months to years). This, in turn, limits the development of clinical rehabilitations of a significant part of the population that chronically experiences abnormal optics in their everyday life. A major experimental challenge comes from the difficulties of empirically isolating neural from optical factors. Technological advances in the field of adaptive optics (AO) offer a unique opportunity to bypass the limits imposed by optical factors while directly assessing visual processing of images free from any optical imperfection (8). AO is a powerful technology that can be used to improve optical systems, including the human eye, by deforming a mirror to correct wavefront distortions of the optics (**Fig.1** and *SI Methods*). AO-correction in “healthy” eyes allows observers to detect a larger range of high-SF information otherwise indiscernible in the presence of typical optical blur, improving visual acuity (9) and contrast sensitivity (10). More importantly, AO correction can be used to assess how changes in optical quality alter neural functions, by making it possible to compare visual functions under fully-corrected optical quality of both typical “healthy” eyes (5,9-11) and those with severe optical abnormalities (12-14).

In this context, keratoconus (KC) represents an ideal model of long-term neural adaptation to optical defects. KC is a severe corneal disease afflicting young to middle-aged, neurotypically-developed adults (**Fig.1a**). In this condition, the corneal stroma progressively thins and assumes a conical shape, resulting in a substantial increase in both lower-order (defocus and astigmatism) and higher-order (HOAs) aberrations. Although HOAs are relatively small in typical eyes, abnormal corneal conditions (e.g., such as KC) can cause large magnitudes of HOAs that cannot be efficiently corrected by conventional optical devices (15). Thus, despite their habitual optical correction, KC patients are chronically exposed to substantially degraded retinal inputs in their everyday life. Prolonged exposure to poor optical quality results in neural adaptation that alters visual processing and limits the benefits of improved optical correction. When tested under full AO-correction, KC patients exhibit considerably poorer visual acuity (13) than that predicted by optical theory or measured in neurotypical (NT) observers with similar AO-corrected optical quality. However, little is known regarding the underlying mechanisms mediating such changes in visual processing following long-term adaptation to severe optical aberrations.

Here, we used state-of-art AO to assess how long-term exposure (i.e., years) to optical defects alters neural processing of visual information in neurotypically-developed humans. To do so, we measured the impact of different amounts of optical aberrations experienced by NT observers and KC patients on the visual system’s ability to detect contrast over a wide range of SFs under AO-corrected optical quality. The visual system is composed of SF-selective filters whose combined sensitivity determines the shape of the contrast sensitivity function (CSF)(16,17). For well over a century, contrast sensitivity has been a valuable tool for measuring the limits of visual perception, as well as for the assessment and early diagnosis of many disorders (17). The detection of alterations in the shape of the CSF can allow inferences about changes in underlying physiological processes, such as in amblyopia (18), autism spectrum disorder (19), and aging (20). As the impact of optical aberrations is more pronounced for high SFs, we expect deficits in contrast sensitivity at high SFs despite correcting all optical aberrations with AO, whereas sensitivity to low-SF information should remain unaffected. However, whether neural processing in KC is simply impaired for visual inputs affected by the eye’s optics or undergoes a more adaptive functional reorganization is unknown. In two experiments, we provide compelling evidence of neural compensation for severely degraded retinal inputs experienced over prolonged periods of time. First, we show evidence of altered CSF and poorer visual acuity in KC patients relative to age-matched NT observers, despite being tested under AO correction. Then, using an equivalent noise paradigm and a computational model of visual processing (21-25), we identify the putative mechanisms underlying changes in visual sensitivity observed in KC patients under AO correction. Overall, our findings reveal that chronic exposure to poor optical quality does not result in neural deficits restricted to high SFs, but rather manifests by large-scale changes in visual sensitivity across a broad range of SFs.

**Figure 1.**
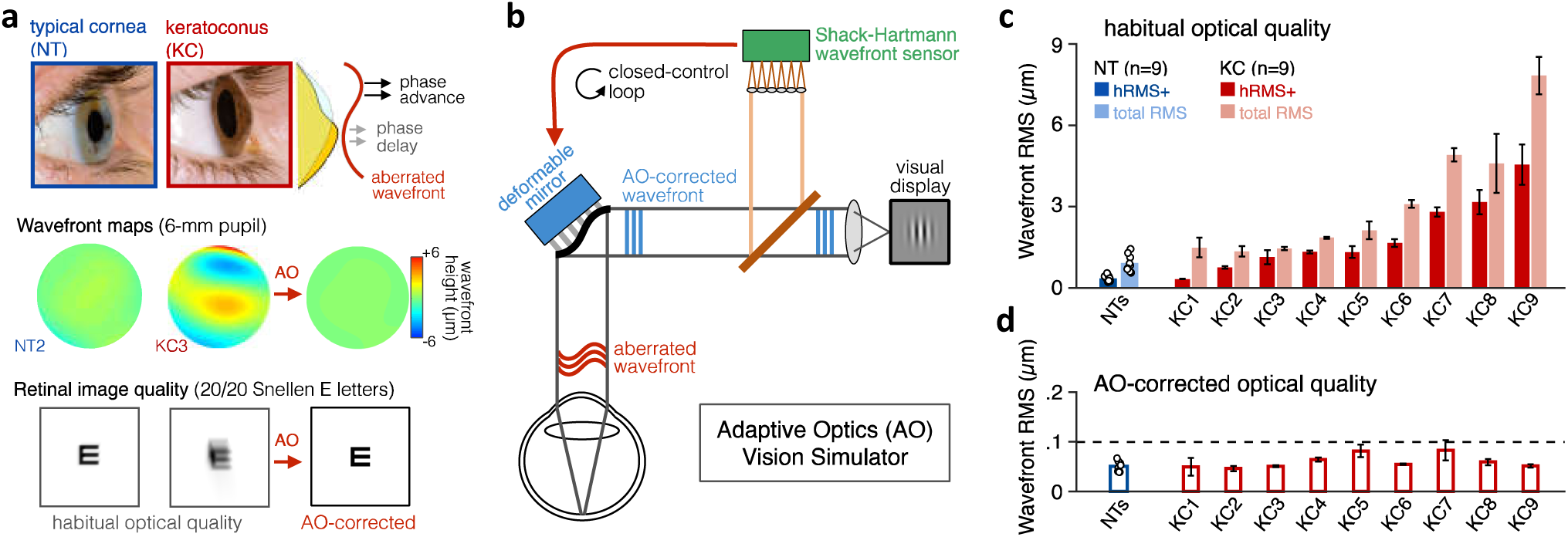
Using adaptive optics to study long-term neural adaptation to poor optics in keratoconus. **(a)** KC is a progressive eye disease affecting neurotypically-developed adults, in which the typically round cornea thins and bulges into a cone-like shape. KC results in large amounts of higher-order optical aberrations (HOAs), as shown on the wavefront maps, which cannot be fully corrected and chronically degrade retinal image quality. Adaptive optics (AO) allows to bypass optical factors and assess visual processing under fully-corrected optical quality. **(b)** The AO vision simulator maintains aberration-free image quality during testing, by measuring the subject’s aberrated wavefront using a Shack-Hartmann wavefront sensor and fully correcting it online with a deformable mirror in closed-control loop. **(c)** Relative to NT eyes, KC eyes were chronically exposed to severe optical defects, with large amounts of HOAs (hRMS+). Group-average RMS and individual data points are presented for NTs. Individual bars for each KC patient show habitual RMS estimates (±1SD). **(d)** AO-correction maintained aberration-free optical quality during visual testing, even in severe KC eyes. Note the large difference in y-axis scale between panels c and d.

## Results

### Habitual aberrations and AO-corrected visual quality

We first needed to quantify the amount of habitual optical aberrations experienced by each observer, before establishing that we can fully correct all optical aberrations to a similar level in both NT observers and KC patients with varying levels of optical defects. Habitual optical quality was estimated using AO (**Fig.1b** and **Fig.S1**) by collecting multiple wavefront measurements for each observer wearing their own, everyday optical corrective method, if any. Wavefront maps were fitted with Zernike polynomials to compute the total RMS and higher-order RMS (hRMS+; HOAs only) for a 6-mm pupil (**Fig.1c**; see also *SI Methods* and **Table S1**). Relative to optical quality in age-matched NT observers (total RMS: 0.91±0.31µm; range: 0.57-1.44), habitual optical quality in KC eyes (total RMS: 3.19±2.2µm; range: 1.35-7.84), remained sub-optimal despite their own optical corrections (*NT-vs-KC total RMS*: Welch’s t-test, t=3.08, *p=0.014*, d=1.45). Specifically, KC eyes were subject to a substantial amount of uncorrected HOAs (hRMS+: 1.90±1.35µm; range: 0.33-4.55), much more than NT eyes (hRMS: 0.35±0.11µm; range: 0.22-0.55) (*NT-vs-KC hRMS+*: Welch’s t-test, t=3.43, *p=0.009*, d=1.62). As previously reported (15), KC eyes were particularly affected by large amounts of vertical coma (mean absolute *Z*_7_ coefficient: 1.38±1.02µm, range: 0.03-2.88), much more than in typical eyes (0.13±0.12µm, range: 0.02-0.34).

Consistent with previous studies from our lab (12-14), our AO system allowed us to measure visual performance while effectively maintaining aberration-free optical quality, even in severe KC eyes (**Fig.1c** and **Fig.S2**). To maximize AO correction during stimulus presentation, observers were trained to blinks between trials and to pause if the perceptual quality got unstable or poor quality during testing (see *SI Methods*). Continuous closed-loop AO correction resulted in a residual wavefront RMS that remained not significantly different from 0.055 µm for both NT eyes (residual RMS: 0.051±.01µm; Student t-test: t(8)=1.24, *p=.251, d=0.41*) and KC eyes (residual RMS: 0.060±.014µm; Student t-test: t=1.13, *p=.292, d=.38*). More important, aberration-free optical quality under AO correction was similar between NT and KC eyes (Student t-test, t=1.64, *p=0.120*, d=0.77), with no relation between AO-corrected RMS and the severity of the habitual RMS (Pearson’s correlation: total RMS r=+.32, *p=.196;* hRMS+ r=+.35, *p=.157*). Thus, online AO correction effectively and continuously corrected all optical aberrations during testing, allowing us to bypass optical factors and assess differences in visual processing between NT and KC observers under similar aberration-free conditions.

### Altered contrast sensitivity function following long-term adaptation to optical defects

In Experiment 1, we assessed the CSF in both KC (N=9) and age-matched NT (N=9) observers under aberration-free conditions. Observers performed an orientation discrimination task at fixation (**Fig.2a**) in which they reported the orientation of ±45º-tilted Gabor patches varying in SF (in cycle per degree or cpd) and contrast. To optimize data collection, we used the *quick CSF* method (18,26), which combines Bayesian adaptive inference with a trial-by-trial information gain strategy to estimate the observer’s CSF as a truncated log-parabola with four parameters (**Fig. 2b** and *Methods*): 1) peak sensitivity, *CS*_max_; 2) peak frequency, *SF*_peak_; > 3) bandwidth, *β*; and 4) low-SF truncation level, *δ*. The low-SF truncation level determines sensitivity at low SFs (*CS*_low_). We also estimated the high-SF cutoff (*SF*_cutoff_) from qCSF fits—a measure of visual acuity. The qCSF method has been used to characterize CSF in both neurotypical and diverse clinical populations (18,26-28).

**Figure 2.**
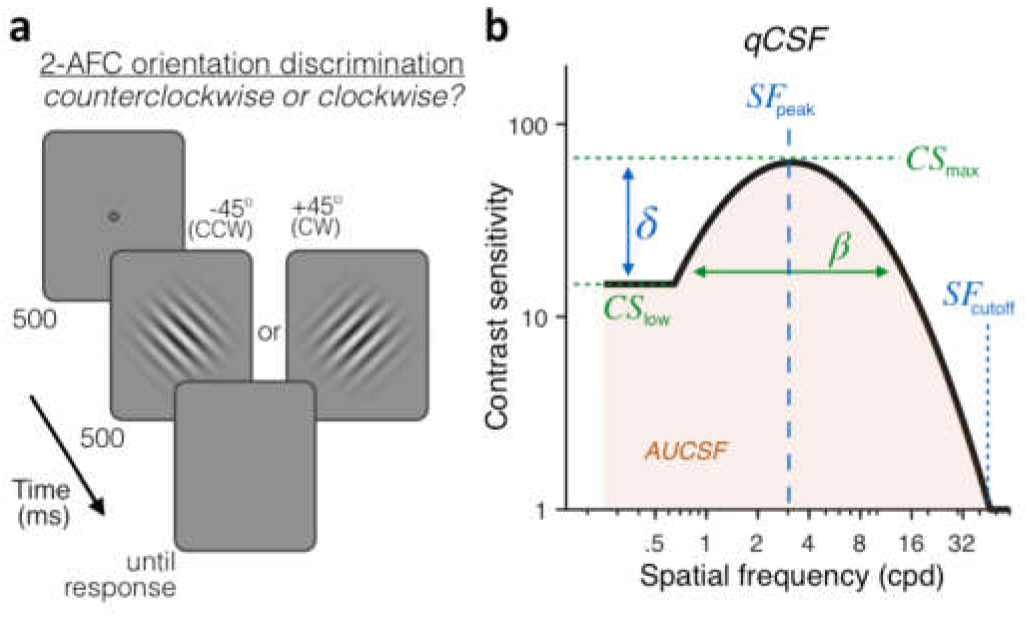
Experiment 1: Contrast Sensitivity Function (CSF) measurement. **(a)** *Stimuli task, and timeline.* Each trial began with a dynamic fixation point. After a blank screen, a ±45°-oriented Gabor stimulus was presented at fixation using a 500ms temporal Gaussian envelope. Stimuli varied in contrast and spatial frequency (SF) from trial to trial. Observers judged whether the stimulus was tilted ±45° from vertical. Full AO correction was maintained during testing. **(b)** *quick CSF method*. The qCSF estimates the CSF through 4 parameters: peak sensitivity (*CS*_max_), peak SF (*SF*_peak_), bandwidth (*β*), and low-SF truncation (*δ*). The area under the CSF (AUCSF), low-SF sensitivity (*CS*_low_), and high-SF cutoff (*SF*_cutoff_) were also estimated from qCSF fits.

Under similar, aberration-free optical quality, KC participants exhibited altered CSF relative to NT observers (**Fig.3a**), showing both impairments for high SFs (*p<0.05*, for SFs≥5.3 cpd) as well as improvements for low SFs (*p<0.05*, for SFs≤0.6 cpd). In other words, even when both groups had matched retinal image quality, there were still substantial differences in visual sensitivity. This pattern of gains and losses changed the shape of the CSF, revealing both impaired sensitivity at high SFs (*SF*_cutoff_; NT: 41.3 cpd [39.2–44.7; 95%-CI]; KC: 32.2 cpd [30.8–35.3]; *p<0.001*) and better sensitivity at low SF (*CS*_low_; NT: 4.92 [4.03–6.46]; KC: 8.64 [6.72–10.22]; *p=0.0012*). Improved low-SF sensitivity reflected reduced low-SF truncation level in KC (*δ*; NT: 1.23 [1.13–1.32]; KC: 0.94 [0.87–1.06]; *p<0.001*). Notably, the impairment in high-SF cutoff correlated with the reduction in low-SF truncation across observers (r= .47, *p=0.047*)—a pattern of results that is suggestive of functional reorganization of visual processing across SFs. The overall result was a shift in peak SF towards lower SFs in KC (*SF*_peak_; NT: 3.42 cpd [3.25– 3.83]; KC: 2.84 cpd [2.67–3.11]; *p<0.001*). No significant changes in amplitude (*CS*_max_; NT: 82.6 [76.2– 94.2]; KC: 75.8 [66.8–88.5]; *p=0.128*) or bandwidth (*β*; NT: 2.83 [2.64–2.98]; KC: 2.77 [2.60–2.92]; *p=0.338*) were observed. These changes resulted in the area under the CSF (AUCSF) to be slightly reduced in KC patients (NT: 2.81 [2.77–2.90]; KC: 2.73 [2.66–2.81]; *p=0.025)*. Note that AUCSF depends on the SF range used. Had we used additional low SFs, where KC patients showed better performance, this difference in AUCSF between groups would have been attenuated, and perhaps eliminated.

**Figure 3.**
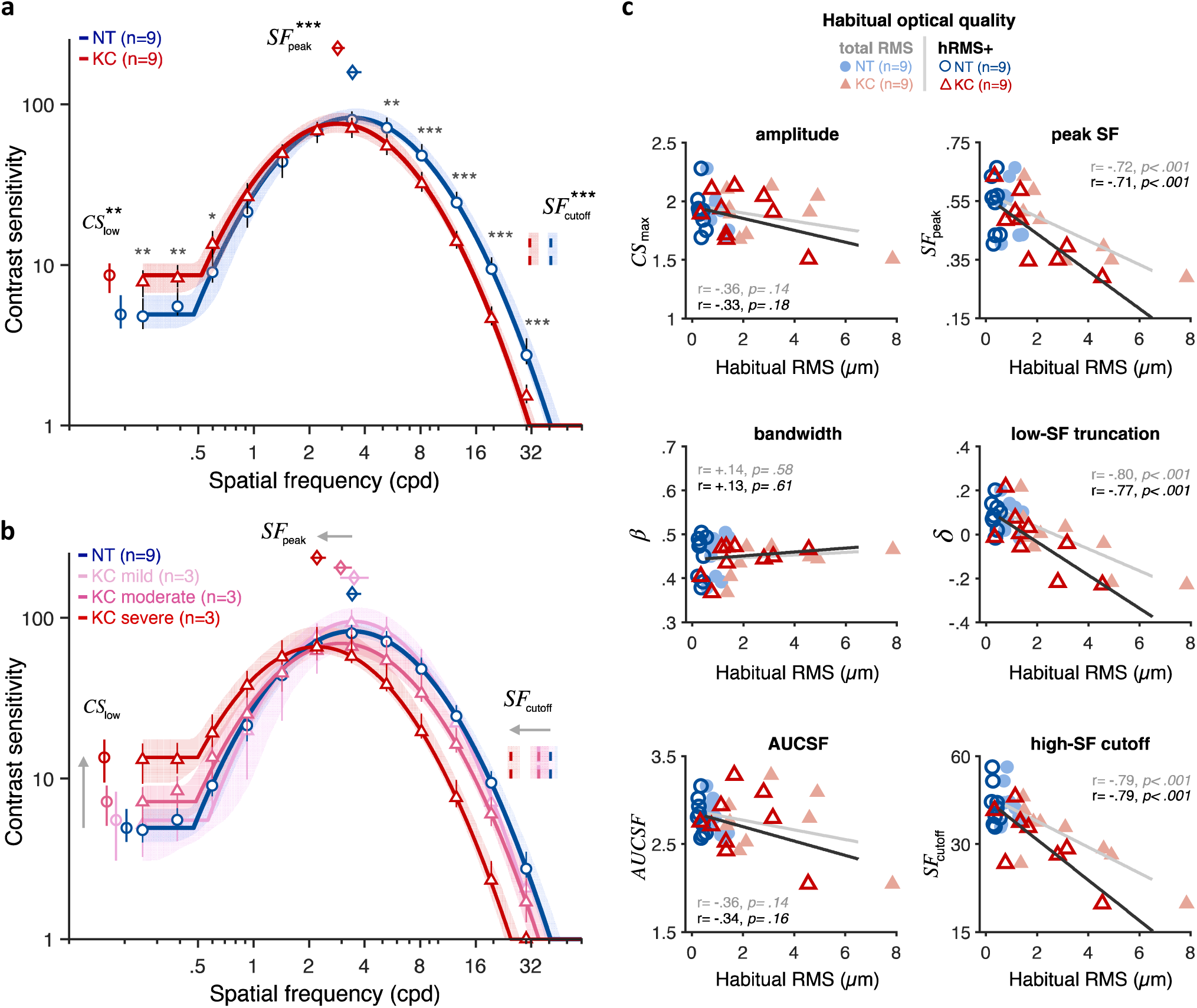
Experiment 1: Altered contrast sensitivity function (CSF) following long-term adaptation to poor optics. All results are for AO-corrected visual images for both keratoconus (KC) and neurotypical (NT) observers. **(a)** *qCSF results*. Relative to NT observers (blue curve), KC patients (red curve) showed altered CSF that were shifted towards lower SFs, with both impaired high-SF sensitivity and improved low-SF sensitivity. Shaded areas and error bars represent bootstrapped 95%-CI. Asterisks indicate significant differences computed from bootstrapping between groups (*: *p<0.05*; **: *p<0.01*; ***: *p<0.001*). **(b)** Same as (a) with KC patients divided into 3 equal groups based on KC severity (see also Fig.S3). **(c)** *Link between CSF and habitual RMS*. Altered CSF under AO correction correlated with the amount of habitual aberrations of each subject. Each panel shows individual parameter estimates plotted as a function of each subject’s habitual RMS (total RMS: light filled symbols; hRMS+: darker open symbols). Linear regression fits and Pearson’s correlation coefficients are plotted for each RMS condition.

Critically, both gains and losses in CSF under aberration-free conditions were more pronounced in severe KC, who experienced large amounts of habitual aberrations in their everyday life (**Fig.3b** and **Fig.S3**). Changes in qCSF parameters under full AO correction correlated with the amount of habitual aberrations each observer experienced in their everyday life (**Fig.3c**; Pearson’s correlation coefficients using total RMS; *SF*_peak_: r=-.71, *p<0.001*; *SF*_cutoff_: r=-.79, *p<0.001*; *δ*: r=-.80, *p<0.001; CS*_low_: r=+.62, *p=0.006*). No correlation was observed for other qCSF parameters (*CS*_max_, *β*, or AUCSF). A similar pattern was observed using hRMS+ as an index of habitual optical quality (**Fig.3c**). Furthermore, visual acuity (VA) measurements using high-contrast Snellen E letters (**Fig.S4** and *SI Methods*) showed poorer acuity in KC patients (15.14±1.64, range: 13.38-17.81) than in NT observers (11.97±0.94, range: 10.58-13.61), despite being tested under full AO correction (*NT-vs-KC logMAR VA*: t=5.34, *p<0.001*, d=2.52). This deficit in high-contrast VA is consistent with previous findings (13) and correlated with the amount of habitual aberrations experienced by each observer (total RMS: r=+.85, *p<.001*; hRMS+: r=+.87, *p<.001*), as well as with the changes in peak SF (r=-.74, *p<.001*), high-SF cutoff (r=-.77, *p<.001*), and notably low-SF truncation (r=-.72, *p<.001*) and low-SF sensitivity (r=+.61, *p=.007*). That is, poorer VA in KC under AO correction was predicted by both impaired high-SF sensitivity and improved low-SF sensitivity. These results indicate that long-term exposure to severe optical aberrations alters the CSF in a way that results in impaired fine spatial vision, but also in better sensitivity to coarser spatial information less affected by optical aberrations. The resulting large-scale change in contrast sensitivity across SFs is expected to optimize visual processing of severely aberrated retinal images caused by KC, but limits perceptual quality under aberration-free optical quality.

### Altered internal noise properties following long-term adaptation to optical defects

To better understand how long-term exposure to optical defects alters visual processing across SFs, we used an equivalent noise paradigm (21-25) where perceptual thresholds are measured as a function of varying external noise levels (see *Methods*). Six participants from each group (**Table S1**) performed an orientation discrimination task under varying levels of dynamic white noise added to the stimuli (**Fig.4a**), judging whether a foveally-presented grating was tilted ±45 from vertical. Stimulus presentation was controlled using the *FAST* method (29), an advanced adaptive psychophysical technique that allowed us to estimate relevant model parameters in just 480 trials per participant (21-25). Changes in perceptual thresholds with external noise result in a characteristic threshold-versus-noise (TvN) function, which can be fitted with the Perceptual Template Model (PTM) (21,22) to quantify the effects of noise on perception (**Fig.4**). The PTM is a computational model of visual processing that had been successfully used to identify the mechanisms underlying perceptual differences for a wide range of brain functions, such as attention (30,31) and perceptual learning (21), as well and between specific populations, such as in amblyopia (32), autism (24), or dyslexia (33). The PTM considers that differences in perceptual performance between groups can result from changes in three possible sources of noise (**Fig. 4b**): (1) internal additive noise (*A*_*add*_); (2) external noise filtering (*A*_*ext*_); and (3) internal multiplicative noise (*A*_*mul*_). Elevated internal additive noise leads to worse perceptual thresholds at low external noise levels, reflecting impaired signal enhancement of the perceptual template (**Fig. 4c**, *upper left*). Poorer external noise filtering impairs performance at higher external noise levels, reflecting poorer signal selectivity (tuning) of the perceptual template (**Fig. 4c**, *upper right*). A combined effect of higher internal additive noise and poorer external noise filtering elevates thresholds at all external noise levels, similarly across difficulty levels (**Fig. 4c**, *lower left*). In contrast, higher internal multiplicative noise (*A*_*mul*_) would also impair thresholds across external noise levels, but via a non-uniform shift in TvN functions across difficulty levels (**Fig. 4c**, *lower right*). Higher *A*_*mul*_ results in stronger elevation in internal noise with higher signal contrast, acting similarly to a contrast gain control mechanism (i.e., compressive response at higher contrasts).

**Figure 4.**
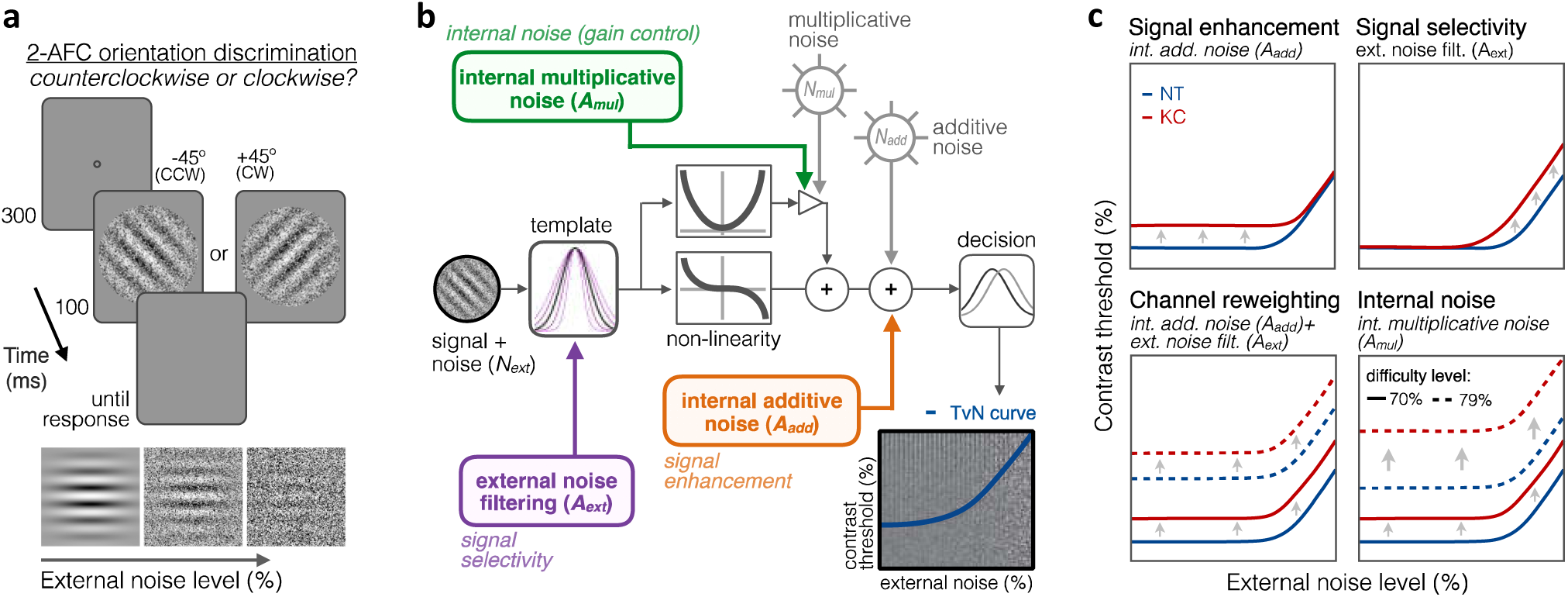
Experiment 2: Equivalent noise paradigm and Perceptual Template Model (PTM). **(a)** *Stimuli, task, and timeline*. In each trial, an oriented grating embedded in dynamic Gaussian pixel noise was presented. Observers judged whether stimuli were tilted ±45° from vertical. **(b)** *Schematic representation of the PTM*. This model consists of five main components: a perceptual template processing signals embedded in external noise (*N*_*ext*_), a nonlinear transducer function, two internal noise sources (multiplicative *N*_*mul*_ and additive *N*_*add*_), and a decision process. The output of the decision process models limitations in sensitivity as equivalent internal noise, with threshold-vs-noise (TvN) curves having a characteristic nonlinear shape. At low external noise, internal noise dominates and added external noise has little effect, resulting in the TvN curve’s flat segment. Once external noise level exceeds that of internal noise, stronger signal contrast is needed to overcome added external noise, resulting in the TvN curve’s rising segment. Adapted from (22). **(c)** *Predictions*. Distinct mechanisms can account for group differences. (1-*upper left*) Elevated internal additive noise (*A*_*add*_) yields increased thresholds at the TvN curve’s flat portion, reflecting signal enhancement changes; (2-*upper right*) Poorer external noise filtering (*A*_*ext*_) limits thresholds at the TvN curve’s rising portion, reflecting signal selectivity changes. (3-*lower left*) A combination of the two result in increased thresholds across external noise levels and difficulty levels, suggesting channel reweighting; (4-*lower right*) Elevated internal multiplicative noise (*A*_*mul*_) would result in a similar pattern across external levels but differences would scale with difficulty levels.

First, we analyzed the data using the conventional PTM (21,22) (see *Methods*) to characterize the relative difference in noise-limiting factors between NT and KC groups. To do so, contrast thresholds were estimated for each observer as a function of external noise, difficulty level, and stimulus SF. Then, we fitted the data with the PTM in a conventional manner using a least-squares procedure (**Fig.5**). Given the large individual variability in KC habitual aberrations and in its impact on the CSF, KC participants were split into 2 groups based on KC severity using a median split (total RMS: 3.85µm; hRMS+: 2.23µm), resulting in three groups based on habitual optical quality (**Table S1**): neurotypical (N=6; total RMS: 0.91±0.33µm; hRMS+: 0.35±0.12µm), mild/moderate KC (N=3; total RMS: 2.19±0.87µm; hRMS+: 1.25±0.46µm), and severe KC (N=3; total RMS: 5.78±1.79µm; hRMS+: 3.51±0.92µm). The data from all SFs were included in the analysis, with the model estimating separate TvNs for each group, each SF, and each difficulty levels (see *Methods*). The resulting TvN curves estimated by the PTM exhibited characteristic nonlinear patterns (**Fig.5**).

**Figure 5.**
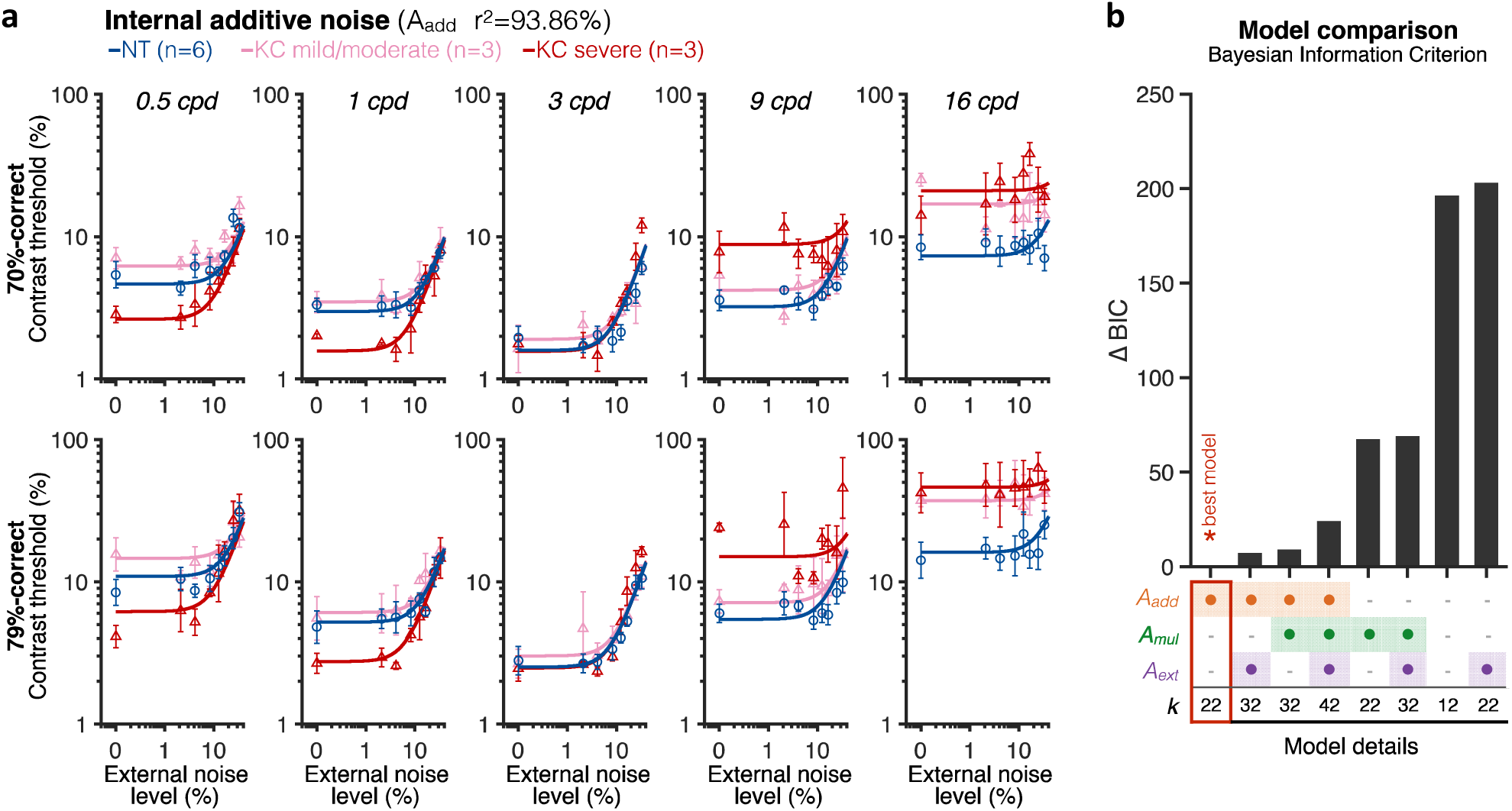
Experiment 2: SF-specific changes in internal additive noise mediate contrast sensitivity differences in KC. **(a)** Contrast thresholds were measured under full AO correction as a function of external noise levels, for five different SFs (0.5, 1, 3, 9, and 16 cpd) and for either 70.71%-correct (*upper row*) and 79.37%-correct (*lower row*) difficulty levels. Relative to NT observers, severe KC patients showed impaired high-SF thresholds, and better thresholds at low SFs. These differences were observed at low, but not at high external noise levels, and were less pronounced in KC patients with mild-to-moderate amounts of habitual optical aberrations. Data were fitted with the PTM to evaluate the contribution of distinct sources of inefficiencies: internal additive noise (*A*_*add*_), internal multiplicative noise (*A*_*mul*_), external noise filtering (*A*_*ext*_), or any combination of these factors. The best model explaining the differences between groups was the internal additive noise model. **(b)** *Model comparisons*. Bayesian Information Criterion (BIC) evidence was computed for each model to identify which model best explained the data while penalizing for greater number of free parameters (*k*). The value of the best model was subtracted from all BIC values (ΔBIC). The best model (ΔBIC=0) was the model assuming solely changes in internal additive noise across SFs between NT and KC groups.

To assess mechanisms underlying differences in contrast sensitivity between groups, several variants of the PTM were fit to the data (**Fig. 5a**; see also **Table S2** and **Fig.S5**): no group difference (null model), changes in internal additive noise (*A*_*add*_), external noise filtering (*A*_*ext*_), or internal multiplicative noise (*A*_*mul*_), mixtures of them (*A*_*add*_+*A*_*mul*_; *A*_*add*_+*A*_*ext*_; *A*_*mul*_+*A*_*ext*_), and a full model assuming changes in all three mechanisms (*A*_*add*_+*A*_*ext*_+*A*_*mul*_). To account for the large differences in the number of free parameters across models, we computed the Bayesian Information Criterion (BIC) evidence that determines which model best fits the data while penalizing for greater number of free parameters. Overall, the differences in contrast thresholds between groups were best explained by a simple model assuming solely SF-dependent changes in internal additive noise (r^2^=93.86; **Fig. 5a**), without changes in external noise filtering and/or internal multiplicative noise. Relative to NT observers, severe KC patients showed a reduction in internal additive noise (*A*_*add*_) by -42.6% and -46.3% at 0.5 and 1 cpd, respectively. Mild/moderate KC patients did not show such benefits at low SFs, but rather an elevation in internal noise by +32.8% and +16.8% at 0.5 and 1 cpd, respectively. At 3 cpd, mild/moderate KC observers were still slightly worse than NT observers due to elevated internal additive noise (+18.9%), whereas severe KC patients showed a negligible reduction in internal noise (−1.6%). At high SFs, all KC participants exhibited impaired contrast thresholds due to large elevations in internal additive noise, by +30.1% and +126.4% at 9 and 16 cpd for mild/moderate KCs, and by +167.7% and +179.7% at 9 and 16 cpd, for severe KC patients. Additional PTM analyses, separately for low and high SFs, further supported SF-specific changes in internal additive noise as the primary mechanism underlying altered CSF following long-term exposure to poor optical quality (**Fig.S6**). The pattern of gains and losses in sensitivity across SFs at low external noise levels was consistent with the altered CSF observed in Experiment 1 (**Fig.S7**).

To better assess individual differences in the underlying sources of altered visual processing across SFs and relate them to each observer’s habitual optical quality, we estimated TvN curves and parameter estimates for each individual participant using a hierarchical Bayesian model (see *Methods*). By assuming that the variability between participants follows a population-level distribution, this method allowed us to estimate group-level “traits” in terms of signal processing given the presence of external noise. We then tested whether the level of internal additive noise, as well as other PTM estimates, correlated with the amount of each participant’s habitual RMS. Consistent with the SF-specific changes in sensitivity observed in severe KC patients, poorer habitual RMS was associated with reduced individual internal additive noise levels at low SFs (0.5 and 1 cpd) (**Fig.6a**; total RMS, r=-.60, *p=0.037*; hRMS+, r=-.58, *p=0.048*). At high SFs, individual noise estimates were more variable, particularly at 16 cpd where we could not reliably estimate the non-linear segment of individual TvN curves. Thus, we restricted our analysis to individual noise estimates from the 9 cpd condition. We found that poorer habitual optical quality was positively correlated with elevated individual internal additive noise levels at high SFs (**Fig.6b**; total RMS, r=+.65, *p=0.022*; hRMS+, r=+.71, *p=0.010*). No correlation was found with either internal multiplicative noise (*A*_*mul*_) or external noise filtering (*A*_*ext*_) for low or high SFs (**Fig.S8**), supporting the fact that the effects of long-term exposure to severe optical aberrations are best explained by an internal additive noise model that assumes SF-specific changes in signal enhancement.

Taken together, the present findings show that long-term exposure to poor optical defects causes a large-scale functional reorganization in visual sensitivity across SFs, which reflects SF-specific changes in signal enhancement mechanisms of SF-selective neurons. The atypical contrast sensitivity and poorer acuity observed in neurotypically-developed adults with severe optical defects, even when all optical aberrations are bypassed using AO correction, reflect long-term neural compensation mechanisms that improve visual processing of the degraded retinal inputs but limit the clinical benefits of improved optical correction in these patients.

**Figure 6.**
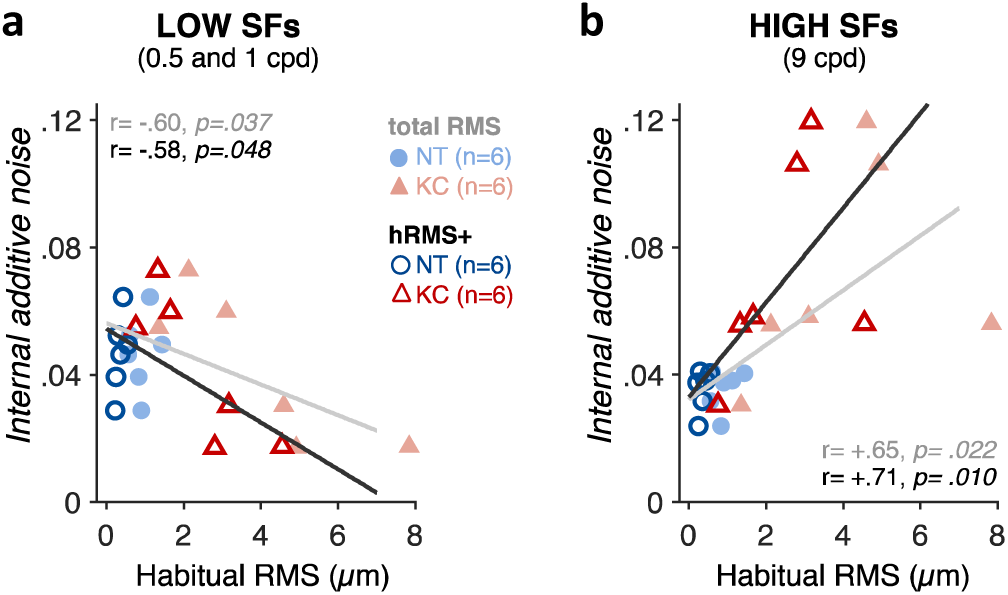
Correlation between individual noise estimates and participants’ habitual optical quality. **(a)** At low SFs (0.5 and 1 cpd), poorer habitual optical quality was associated with reduced internal additive noise. **(b)** At high SFs (9 cpd), the opposite pattern was found, with poorer habitual optical quality being associated with elevated internal additive noise. Other noise estimates (i.e., internal multiplicative noise and external noise filtering) did not correlate with habitual optical quality (see Fig.S8).

## Discussion

The present study combined advanced optics, psychophysical and computational methods to assess neural adaptation in response to long-term (i.e., up to 20 years) exposure to poor optical quality. An AO vision simulator was used to fully bypass optical limitations and directly access and isolate visual processing in human adults, who are otherwise exposed to various amounts of optical aberrations. Our results provide evidence of a large-scale functional reorganization of sensory resources following long-term exposure to severely degraded retinal inputs. Under fully-corrected optical conditions, KC patients showed a loss in sensitivity to high-SF information, which was due to poorer signal enhancement mechanisms of high-SF selective filters. Moreover, severe KC patients showed enhanced sensitivity to low-SF information, which reflected improved signal enhancement of low-SF selective filters. Notably, magnitudes of observed low-SF enhancements and high-SF impairments were correlated across observers, and both effects were more pronounced in more severe KC cases. This pattern of gains and losses in visual sensitivity reveals fundamental properties of adaptive neural mechanisms compensating for the chronic exposure to blurred retinal images.

A key function of sensory systems is to adapt to our sensory environment. Contrast sensitivity represents the foundation for the brain’s coding of visual information and is central for characterizing both visual function and clinical disorders. The shape of the CSF likely reflects the visual system’s sensitivity to stimulus properties that are useful for perception. In neurotypical observers, the drop in sensitivity at high SFs is mostly due to blurring from the eye’s optics (17,34). Optical blur reduces the strength and reliability of high-SF retinal inputs, leading to weak and unreliable neural responses to fine spatial details. In KC, optical aberrations become ∼6 times worse compared to typical levels (15), strongly attenuating high-SF retinal signals and depriving the visual system from fine spatial information. Our findings show that the severe and prolonged exposure to optically degraded retinal inputs under natural viewing results in neural insensitivity to fine spatial details, even when viewing images that are near-perfect optically. Using the PTM, we demonstrated that this neural insensitivity to high-SF signals was due to elevated internal additive noise levels (i.e., reduced signal enhancement) of high-SF filters. Internal noise impairs the reliability of sensory representations, and is a key factor in perceptual variability within the nervous system (35). This increase in internal noise levels of high-SF filters can account for the fact that when tested under AO correction, KC patients showed considerably poorer high-contrast letter acuity than observers with typical optical quality whose AO-corrected acuity approaches the limit imposed by photoreceptor sampling (13,14).

More important, chronic exposure to degraded high-SF inputs did not solely induce impairments in high-SF sensitivity under AO correction. Participants with severe KC also showed better contrast sensitivity to low-SF signals, which was mediated by reduced internal additive noise levels (i.e., improved signal enhancement) of low-SF filters. Thus, the effects of long-term exposure to poor ocular optics cannot be simply characterized as deficits at high SFs, but rather as altered visual sensitivity across SFs via changes in signal enhancement mechanisms that attenuate high-SF signals while enhancing low-SF information. This pattern of gains and losses in sensitivity can be understood by considering how habitual optics of KC eyes affect retinal image quality. While optical blur strongly degrades high-SF signals, lower SFs are less compromised by blur and become predominant and more reliable than higher SFs, especially under large amounts of blur. As a result, KC patients are in a low-pass state of adaptation during natural vision, relying more on low-SF information to interact with their environment. Following short-term blur adaptation, humans with typical optical quality usually show enhanced sensitivity for high SFs and reduced sensitivity for low SFs (36,37). In contrast, our results show that blur adaptation over considerably longer periods of time has opposite effects on contrast sensitivity, supporting the notion that distinct neural adaptation mechanisms operate over different timescales (6,7).

Adaptation acts to maintain sensory systems operating within the limited dynamic range afforded by the brain’s limited resources (38), serving as optimal resource allocation that can manifest in large-scale reorganization in visual sensitivity (39). In this context, the CSF reflects an optimal allocation of the brain’s limited sensory resources to process a wide range of spatial details. The pattern of gains and losses in sensitivity we observed could reflect neural compensation mechanisms that optimize visual processing to the structure of the degraded retinal inputs. For instance, short-term adaptation to different stimulus speeds results in a large-scale reorganization of the spatiotemporal CSF, which optimizes visual sensitivity to the new environment (39). Physiologically, cells in the primary visual cortex (V1) adaptively change their responses to match the input properties of natural (broadband) stimulation and enhance information transmission in visual processing (40). What is unknown, however, is what kinds of neural changes occur in fully-developed visual systems during years of adaptation periods, as experienced by patients with severe optical defects such as KC patients. We conjecture that long-term exposure to severely degraded retinal image quality results in a reallocation of the brain’s limited sensory resources, improving visual processing in KC patients across many stimuli and tasks. Indeed, while KC participants suffer from poorer visual acuity under full AO-correction, they actually show better acuity (i.e., by +24%) under their own optical quality than typical observers tested under the same degraded optical conditions (41).

Our methods are largely agnostic about the neural bases of long-term adaptation to blur. We hypothesize that our findings are likely to be due, at least partially, to neural changes occurring in V1. Pooled responses of V1 neurons can predict behavioral performance in contrast sensitivity tasks (42), and the perceptual CSF and neuronal CSF in V1 are highly correlated (43), sharing a similar shape, amplitude, and peak SF. That is, the shape of the perceptual CSF is assumed to reflect properties of V1 neurons selective to a wide SF range. While responses in the lateral geniculate nucleus (LGN) are primarily low-pass in SF, the majority of V1 neurons possess bandpass characteristics. The fall-off of the CSF at low SFs is generally attributed to lateral inhibition in retinal ganglion cells (44), as well as normalization–a canonical neural computation in which the response of an individual neuron is divided by the summed activity of a pool (population) of neurons (45). As retinal mechanisms tend to be less plastic, long-term adaptation effects on visual sensitivity observed in KC patients could reflect changes in normalization in V1. Divisive normalization plays an important role in determining visual sensitivity, serving as a form of homeostatic control of V1 functional properties that optimize the network nonlinearities to the statistical structure of the visual input (46). Here, severe optical blurring in KC eyes would reduce the contribution of high-SF neurons to the normalization pool, in turn increasing the response of neurons selective to lower SFs. Another, not mutually exclusive, neural account could involve interactions between SF-selective channels (37,47,48). SF-specific inhibitory mechanisms have been shown to refine SF selectivity of V1 neurons (47). SF-specific inhibition of low SFs plays a key role in the generation of bandpass V1 selectivity (48), shifting preferred spatial selectivity to higher SFs. Moreover, adaptation effects at different spatial scales are not independent (37), ruling out simple linear filter models of cortical processing. Changes in normalization and/or SF-specific inhibition mechanisms would involve large-scale changes in neural sensitivity across many V1 neurons selective to a wide range of SFs, consistent with the SF-specific changes in sensitivity we observed in KC patients. Note that we cannot unequivocally rule out other types of plasticity that could affect the shape of the CSF, such as changes in the number of neurons selective for different SFs (16).

The effects of prolonged exposure to poor optical quality may also be interpreted as a form of perceptual learning. Under natural viewing, KC patients who are chronically exposed to severely degraded images have to relearn how to interact with their environment, which is extremely rich in terms of tasks and stimulus properties. Visual adaptation (2) and perceptual learning (49) are two forms of experience-dependent plasticity that can interact, despite having distinct neural mechanisms and perceptual consequences. For instance, perceptual learning has been shown to reconfigure the effects of visual adaptation (50): While adaptation reduced sensitivity before training, learning while in an adapted state reversed the effects of adaptation, leading to an overall benefit following training (50). This reversal in adaptation effects following repetitive training was specific to the trained, adapted state, while untrained adapted states were associated with significant costs in sensitivity after training (50). Such interactions between sensory adaptation and perceptual learning mechanisms may also account for some of the differences in adaptation effects over brief and long-term timescales (6,7). Long-term adaptation to severe optical aberrations induces long-lasting effects on visual processing that limit the clinical benefits of improved optical correction. Our findings identified specific mechanisms at play that should be considered in the clinical treatment of patients with optical defects. For instance, re-adaptation under improved optical correction devices could be combined with targeted perceptual learning paradigm (14) to enhance the speed, efficiency and generality of neural rehabilitation in patients with altered neural processing due to chronic exposure to poor optical quality.

In summary, the present study furthers our understanding of the impact of long-term (years) exposure to severely degraded habitual optics on visual sensitivity to a wide range of SFs. Our findings support the presence of marked neural plasticity in the adult visual system, which allows the brain to longitudinally respond to optically-related sensory loss. Using AO correction, we were able to bypass optical factors in KC patients and uncovered a large-scale functional reorganization of visual sensitivity mediated by SF-specific changes in signal enhancement mechanisms. The CSF provides vital insights into fundamental properties of human vision, especially in individuals with spatial vision deficits (18). Our results show that optical correction alone is unlikely to fully improve patients’ perceptual quality. Instead, clinical rehabilitation approaches should take into consideration changes in visual sensitivity resulting from chronic exposure to poor optics. An important follow-up question in this context will be to assess how fast and to what extent the visual system of KC-afflicted individuals can re-adapt to perfectly-corrected retinal images.

## Methods

For details, see *SI Methods*

### Participants

Nine keratoconus (KC1-9) and nine neurotypical (NT1-9) observers participated in this study (see **Table S1** for demographic information). The Research Subjects Review Board at the University of Rochester approved all experimental protocols, and informed written consent was obtained for all participants. Further information is provided in *SI Methods*.

### Adaptive Optics Vision Simulator (AOVS)

The AOVS allowed us to bypass optical factors during psychophysical testing by measuring and correcting all optical aberrations (**Fig.1b** and **Fig.S1**). The AOVS consisted of a custom-built Shack–Hartmann wavefront sensor and a deformable mirror, to measure and correct the subject’s wavefront aberrations, respectively. A calibrated visual display was used for psychophysical measurements, which were performed at fovea under white light conditions. The AOVS was operated in continuous closed loop (∼8 Hz) to fully correct all aberrations over a 6-mm pupil during testing (**Fig.S2**). Wavefront measurements were collected for each observer using their everyday correction method, if any, and fitted to individual Zernike polynomials to estimate the habitual RMS error (total RMS or hRMS+; **Fig.1c**), reported in microns (*µm*). As detailed and demonstrated previously (11,13,14), our AOVS provided stable, aberration-free optical quality during testing, in both typical and severely aberrated eyes (**Fig.1d** and **Fig.S2**). Further details are provided in *SI Methods*.

### qCSF measurements

The contrast sensitivity function (CSF) of each subject was measured using a 2-AFC orientation discrimination task (**Fig.2a**). In each trial, a Gabor stimulus (Gaussian envelope SD: 0.75 dva) oriented ±45 was presented at fixation using a 500-ms temporal Gaussian ramp. Observers reported whether the stimulus was tilted clockwise or counterclockwise, with auditory feedback for both correct and incorrect responses. The *qCSF* method (18,26) was used to estimate 81%-correct contrast thresholds over a broad SF range (0.25-30 cpd, with 12 equal log-step values). The *qCSF* describes the CSF as a *truncated log-parabola* with four parameters (**Fig.2b**): 1) peak sensitivity, *CS*_max_; 2) peak frequency, *SF*_peak_; 3) bandwidth, *β* (full width at half maximum); and 4) low-SF truncation level, *δ*. Moreover, contrast sensitivity at low SF (*CS*_low_) and high-SF cutoff (*SF*_cutoff_) were also derived from qCSF fits. Confidence intervals (95%-CIs) and *p*-values were computed from bootstrapping. Further dtails are provided in *SI Methods*.

### Equivalent noise paradigm

Contrast thresholds were measured using a 2-AFC orientation discrimination task as a function of external noise levels (**Fig.4a**). In each trial, a ±45º oriented grating (cosine-envelope diameter: 2 dva) embedded in dynamic white noise (8 levels, 0-.33 SD) was presented for 100ms at fixation. Observers reported the target orientation on each trial, with auditory feedback for both correct and incorrect responses. Stimulus presentation was controlled using the *FAST* method (29) to estimate contrast thresholds at each external noise level, for both 70.71%- and 79.37%-correct difficulty levels. Five SFs (0.5, 1, 3, 9 and 16 cpd) were tested across different experimental sessions. Similar to Park and colleagues (24), data were pooled from the *FAST* structures to estimate psychophysical thresholds using a Bayesian model fitting method implementing a Markov Chain Monte Carlo (MCMC). Further details are provided in *SI Methods*.

### Perceptual Template Model (PTM)

Contrast thresholds for two difficulty levels (70.71% and 79.37%) and across external noise levels were fitted with the PTM (21,22). The PTM consists of five main components (**Fig.4b**): a perceptual template tuned to the signal, a non-linear transducer function, an additive (*N*_*add*_) and multiplicative (*N*_*mul*_) internal noise sources, and a decision process that determines contrast threshold at a specific difficulty level (*d’*) as:

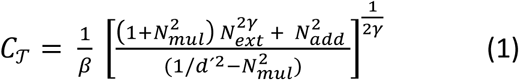

where the input (signal + external noise *N*_*ext*_) is filtered through a perceptual template, resulting in signal enhancement via a gain factor *β*. The output is then transformed through a nonlinear transducer function that amplifies inputs to the *γ*^*th*^ power. Both internal additive noise (*N*_*add*_) and internal multiplicative noise (*N*_*mul*_) sources are added to the output. *N*_*mul*_ remains constant across signal levels, while *N*_*add*_ is proportional to the signal strength. *N*_*ext*_ is manipulated by the experimenter along with the input signal. The PTM considers that performance differences between groups result from changes in three possible sources of inefficiency (**Fig.4b,c**): 1) internal additive noise (*A*_*add*_), reflecting changes in signal enhancement of the perceptual template; 2) external noise filtering (*A*_*ext*_), corresponding to changes in signal selectivity of the perceptual template; 3) changes in internal multiplicative noise (*A*_*mul*_), acting as a contrast gain control mechanism compressing the perceptual template’s responses to signal contrast. Group averages of independently estimated thresholds were used to fit the conventional PTM across SFs. To characterize group differences between NT and KC observers at each SF, the PTM introduces three coefficient indices (*A*_*mul*_(SF,group), *A*_*add*_(SF,group), and *A*_*ext*_(SF,group)) to Equation (1):

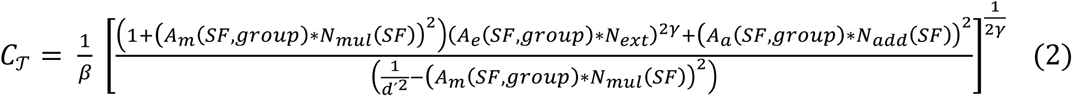

*N*_*add*_(SF) and *N*_*mul*_(SF) varied across SFs to reflect sensitivity differences across SFs. Coefficient indices for NT were fixed to 1. We compared the NT group to 2 groups of KC participants (mild/moderate and severe). To estimate whether and how the three types of noise could account for group differences, we compared the results obtained by fitting the eight possible models (**Fig.S5** and **Table S2**), ranging from no change in any of the noise-reduction mechanisms to the full model with changes in all three noise-reduction mechanisms. Given the large differences in the number of free parameters across models, we computed the Bayesian Information Criterion (BIC) evidence for model selection. The BIC selects the best model while correcting for overfitting by introducing a penalty term for the number of free parameters in the model. Further information is provided in *SI Methods* and *SI Results*.

### Hierarchical Bayesian PTM analysis

To better link habitual optical quality and individual variability in PTM noise estimates, the PTM was fit to each observer using a hierarchical Bayesian modeling technique. This technique estimates PTM parameters for each participant within a population, providing a better understanding of individual variability. For each participant, we estimated three parameters *N*_*mul*,_ *N*_*add*,_ and *W*_*ext*_. We assumed a fixed *β* (1.25) and *γ* (2) for all participants to simplify the model, which was within a reasonable range reported in previous studies (21,22,24). An MCMC technique was used to sample from the posterior distributions and estimate the free parameters. Further information is provided in *SI Methods* and *SI Results*.

## Author contributions

A.B., K.R.H., D.T. and G.Y. designed the study. A.B. and G.Y. collected the data. A.B., W.J.P., R.Y.Z., D.T. and G.Y. analyzed the data. A.B., D.T. and G.Y. wrote the manuscript, and all authors commented on it.

## Acknowledgements

This work was supported by NIH/NEI grant EY014999, Research to Prevent Blindness, and Schmitt Program on Integrative Neuroscience from the University of Rochester Medical Center. We thank Dr. Tara C. Vaz, O.D. and Dr. Len Zheleznyak for their help with participant screening and testing, and Olga Pikul for her help with patient recruitment.

