## Supplemental materials for "Functional reorganization of sensory processing following long-term neural adaptation to optical defects"

### SI Methods

***Participants.*** A total of 9 keratoconus observers (KC1-9; mean age: 37±13, range: 24-55) and 9 neurotypical observers (NT1-9; mean age: 29±7, range: 21-45) participated in this study (see **Table S1** for demographic information). Participants were cyclopleged with tropicamide (1%) to dilate the pupil and paralyze accommodation during the experiment. All participants were screened prior to the study by one of our ophthalmologists, providing standard information (e.g., corneal curvature, refractive error) and ensuring that dilation was safe. An additional screening session was performed to ensure we could obtain good quality wavefront measurements and reach stable aberration-free condition under AO correction. Three potential KC participants did not pass this second screening stage and were not tested further along with KC1-9. The Research Subjects Review Board at the University of Rochester Medical Center approved all experimental protocols. Informed written consent was obtained from all participants prior to participation. Participants were compensated $12/hour.

**Table S1.** Participant information (KC: keratoconus; NT: neurotypical; AO: adaptive optics)

| Participant | Gender | Age (years) | total RMS (μm) 6-mm pupil | hRMS+ (μm) 6-mm pupil | AO RMS (μm) 6-mm pupil | Experiment | Habitual RMS severity |
|---|---|---|---|---|---|---|---|
| KC1 | F | 39 | 1.49±0.36 | 0.33±0.01 | .050±.018 | 1 | mild |
| KC2 | F | 24 | 1.35±0.19 | 0.76±0.05 | .046±.005 | 1, 2 | mild |
| KC3 | M | 26 | 1.46±0.06 | 1.14±0.26 | .051±.002 | 1 | mild |
| KC4 | M | 46 | 1.85±0.04 | 1.33±0.05 | .064±.004 | 1 | moderate |
| KC5 | M | 27 | 2.12±0.33 | 1.33±0.22 | .082±.013 | 1, 2 | moderate |
| KC6 | M | 28 | 3.10±0.14 | 1.66±0.14 | .055±.001 | 1, 2 | moderate |
| KC7 | M | 28 | 4.92±0.24 | 2.80±0.17 | .083±.020 | 1, 2 | severe |
| KC8 | M | 54 | 4.60±1.08 | 3.16±0.45 | .059±.006 | 1, 2 | severe |
| KC9 | M | 55 | 7.84±0.69 | 4.55±0.75 | .051±.003 | 1, 2 | severe |
| KCs (N=9) | 7/9 M | 37.3±13 | 3.19±2.20 | 1.90±1.35 | .061±.01 | - | |
| NTs (N=9) | 9/9 M | 29±7.4 | 0.91±0.31 | 0.35±0.11 | .051±.01 | - | |

***Adaptive Optics Vision Simulator.*** An adaptive optics vision simulator (AOVS) allowed us to bypass any optical factors during psychophysical testing by measuring and correcting all monochromatic and polychromatic aberrations. As illustrated in a simplified schematic (**Fig.1b**; see also **Fig.S1**), the AOVS consisted of a custom-built Shack–Hartmann wavefront sensor to measure the wavefront aberrations, a deformable mirror (ALPAO DM97, Montbonnot, France) to correct subjects' wavefront aberrations, an artificial pupil set to 5.8 mm, and a calibrated visual display for psychophysical measurements. Wavefront aberrations were measured for a 6-mm pupil at fovea using an infrared (840±20 nm) superluminescent diode. The AOVS was operated in continuous closed loop (~8 Hz), allowing to fully correct all aberrations over a 6-mm pupil using the deformable mirror. Visual testing was performed at fovea under white light conditions using a modified digital light processor display (Sharp XR-10X, Abeno-ku, Osaka, Japan) operating at 8 bits with 1024x768 (75 Hz) resolution, sustending 3.56º X 2.67º degrees of visual angle (dva). The display was calibrated with a PR-650 SpectraScan Colorimeter (Photo Research, Chatsworth, CA) and luminance precision was increased to 10.7 bits using the bitstealing technique. A dental impression bite bar mounted to motorized translation stages (x, y, z) with adjustable lateral headrests was used to stabilize head position and maintain pupil alignment during visual testing (**Fig.S1**).

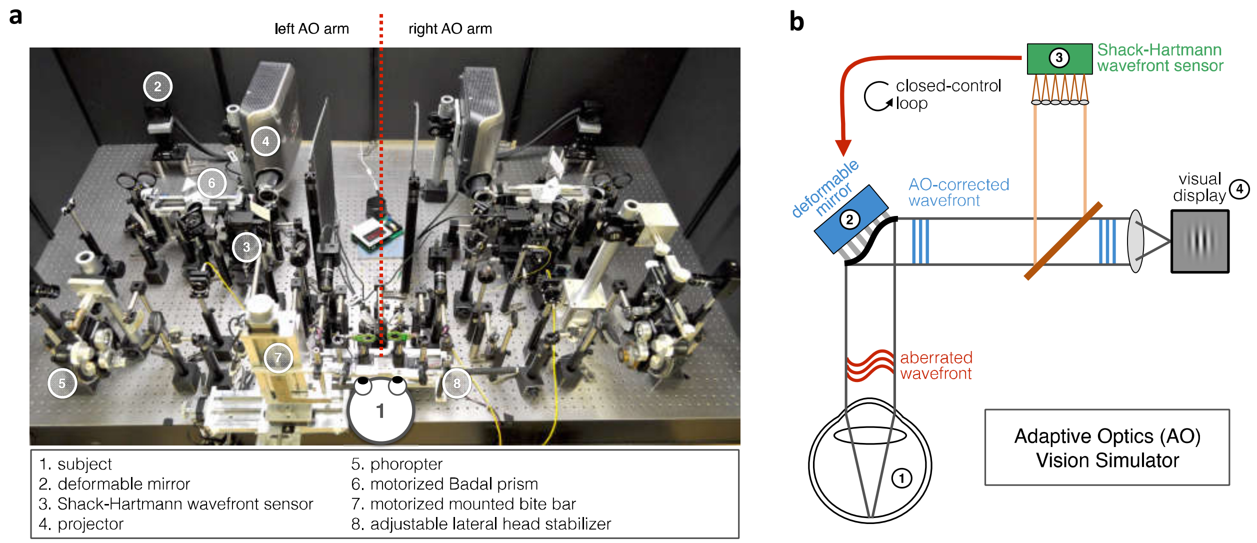


**Figure S1.** *Adaptive Optics Vision Simulator (AOVS)*. **(a)** Picture of the AOVS used in the Yoon's lab at the University of Rochester Medical Center. **(b)** Schematic description of the AOVS. The AOVS allows to measure the aberrated wavefront of the subject's eye using a Shack-Hartmann wavefront sensor, and fully correct it online with a deformable mirror in closed-control loop to ensure aberration-free image quality.

***Optical quality estimation.*** Wavefront measurements were collected for each observer using their everyday correction method, if any, allowing us to estimate each participant's habitual optical quality (**Fig.1c**). Wavefront aberrations were fitted to individual Zernike polynomials up to the 10$^{th}$ order, with 65 Zernikes coefficients. The square root of the sum of Zernikes coefficients aberrations was used to estimate the overall RMS error (total RMS: all Zernikes coefficients) and higher-order aberration (HOAs) RMS error (hRMS+: 6-65$^{th}$ Zernikes coefficients), reported in microns ($\mu m$). Wavefront measurements were also collected during visual testing to estimate both the quality and stability of the AO-correction (**Fig.1d**). Full correction of all Zernikes coefficients was applied for the entire duration of each testing session, with the exception of defocus for which appropriate defocus values were used to correct axial chromatic aberrations. Subjective defocus values were determined for all participants by asking them to adjust defocus using a motorized Badal prism to make a high-contrast Snellen letter "E" as sharp as possible, while correcting all other aberrations using AO. Then, through-focus high-contrast visual acuity measurements were used to objectively verify that the subjective defocus value applied during AO-correction resulted in a very focused, perceived image and maximal VA performance. After this initial session, each experimental session started with VA measurements to ensure the quality and stability of the AO correction before data collection. To maximize AO correction during stimulus presentation, participants were trained to blinks between trials and to stop if the perceptual quality got unstable or poor quality. To minimize the influence of blinks when estimating the residual RMS under AO correction, the median RMS was computed across time for each measurement (**Fig.S2**). Then, the average residual RMS was computed for each participant from multiple wavefront measurements collected under AO correction (**Fig.1d**). As detailed and demonstrated in previous work (11,13,14), our AOVS provided stable, aberration-free optical quality during visual testing, in both typical and severely aberrated eyes.

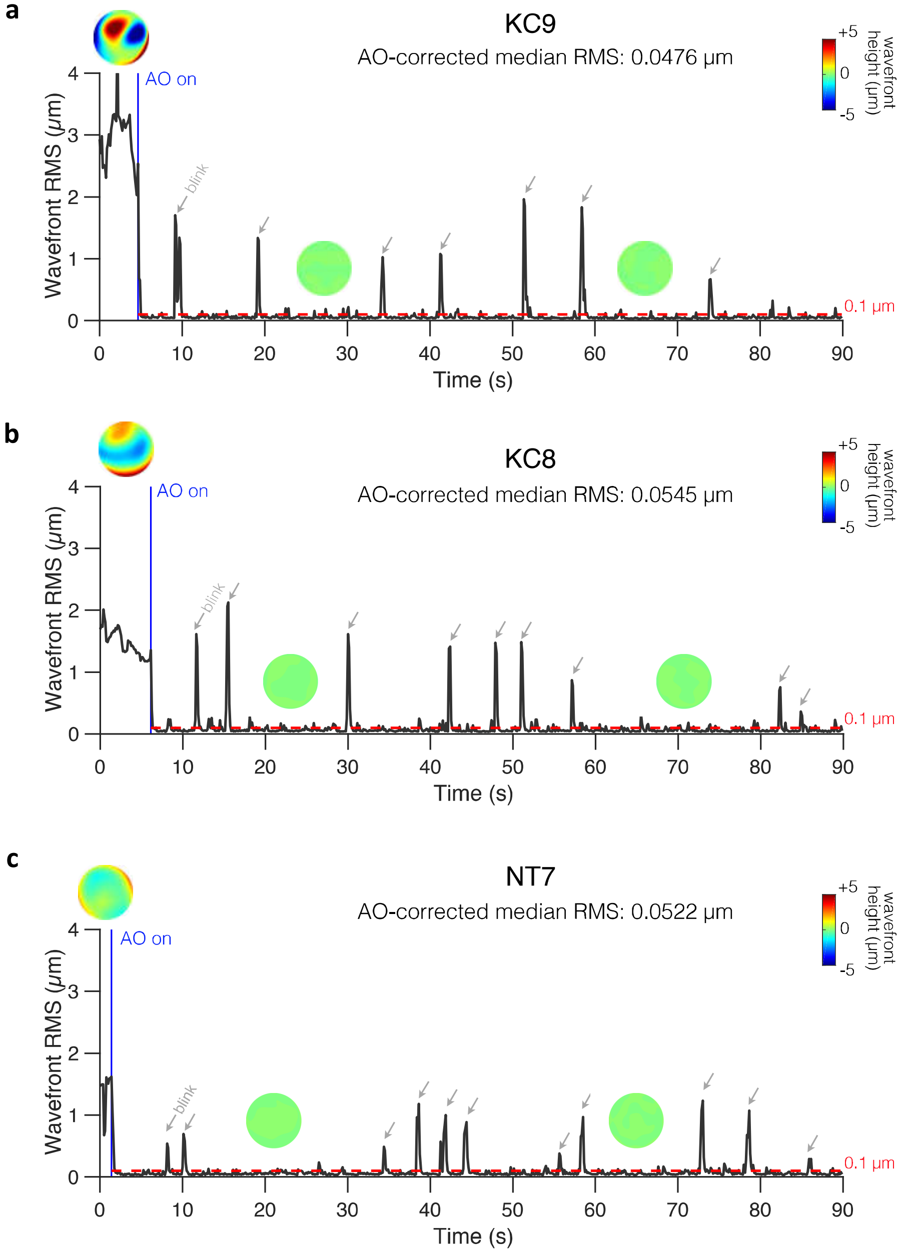


**Figure S2.** *Examples of online AO correction in typical and keratoconus eyes.* **(a)** KC9. **(b)** KC8. **(c)** NT7. Closed-loop AO correction (AO on; blue vertical line) corrected all aberrations and maintained residual total RMS significantly below 0.1 µm during visual testing (red horizontal dashed line). To maximize AO correction during stimulus presentation, participants were trained to blinks between trials and to stop if the perceptual quality got unstable or poor quality. Spikes correspond to blinks (indicated by gray arrows) during which AO correction was paused. The median RMS during AO correction was computed for each measurement to minimize the influence of blinks when estimating the residual RMS under AO correction. To improve AO-correction performance, severe KC eyes were fitted with a scleral lens right by one of the ophthalmologists at the University of Rochester Medical Center before visual testing. Thus, the wavefront patterns before AO correction in panels (a,b) do not reflect the magnitude and pattern of habitual optical aberrations experienced by these KC patients. Note the difference in total RMS before AO correction between NT (c) and KC (a,b) observers. See also Movie S1 from (a).

***Visual acuity measurements.*** Visual acuity (VA) thresholds were measured using a 4 alternative forced choice letter orientation task (**Fig.S4**), in which observers were judged whether a high-contrast Snellen "E" letter presented for 250 ms at fixation was oriented upward, downward, leftward or rightward. Stimuli were black letters presented on a white background. The size of each Snellen E letter was adjusted from trial-to-trial using a staircase method to estimate 62.5%-correct VA thresholds (in logMAR), with 40 trials per staircase. Multiple VA thresholds were collected under AO correction for each participant. Each experimental session also started with VA testing to ensure the quality and stability of the AO correction before starting data collection.

***Contrast sensitivity measurements.*** The contrast sensitivity function (CSF) of each participant was measured using a 2-AFC orientation discrimination task (**Fig.2a**). Each trial began with a dynamic fixation point. After a blank screen, a Gabor stimulus (Gaussian envelope SD: 0.75 dva; SF range: 0.25-30 cpd with 12 equal log-step values) oriented ±45° was presented at fovea. A 500-ms temporal Gaussian ramp was used to blend stimuli into the background and avoid onset/offset transients, and a brief tone signaled stimulus onset to reduce temporal uncertainty. Participants were asked to report whether the stimulus was tilted clockwise or counterclockwise. Auditory feedback was provided for both correct and incorrect responses. We used the *qCSF* method (18,26) to estimate 81%-correct contrast thresholds over a broad SF range. The *qCSF* method is a Bayesian adaptive strategy using *a priori* knowledge about the CSF's general form to obtain reasonably accurate estimates of sensitivity across SFs with as little as 50-100 trials. The *qCSF* method describes the CSF as a *truncated log-parabola* with four parameters (**Fig.2b**): 1) the peak sensitivity (amplitude) $CS_{\text{max}}$; 2) the peak frequency $SF_{\text{peak}}$; 3) the bandwidth $\beta$ (full width in octaves at half maximum); and 4) the truncation level at low SF $\delta$. Without truncation, the CSF is defined as a function of the stimulus frequency ($f$) in decimal log as a *log-parabola CS´(f)*:

$$CS'(f) = \log_{10}(CS_{\text{max}}) - \log_{10}(2)\left(\frac{\log_{10}(f) - \log_{10}(SF_{\text{peak}})}{\log_{10}(2\beta)/2}\right)^2 \qquad (1)$$

This *log-parabola* is then truncated at SF below the peak SF with the truncation parameter $\delta$, which determines contrast sensitivity at low SF ($CS_{\text{low}}$):

$$CS(f) = CS'(f)\,, \qquad sf \geq SF_{\text{max}}, \qquad (2)$$

$$CS_{\text{low}}(f) = \log_{10}(CS_{\text{max}}) - \delta\,, \qquad sf < SF_{\text{max}} \text{ and } SC'(f) < CS_{\text{max}} - \delta$$

In addition, the high-SF cutoff ($SF_{\text{cutoff}}$) was estimated from the qCSF fits, corresponding to the frequency at which $CS(SF_{\text{cutoff}}) = 0$ (i.e., 100% contrast). The test procedure was similar to that described in previous studies (18,26). Briefly, the stimulus space consisted of gratings with contrasts ranging from 0.1% to 99% in steps of 1.5 dB and SF from 0.25 to 30 cycles per degree (cpd). After familiarization with the task, participants performed around 6-7 qCSF runs of 100-trials each (mean number of runs: 6.8±2.6 for NTs, 7±2.8 for KCs), which were then combined to compute individual qCSF functions using all trials. Confidence intervals (95%-CIs) and p-values were computed from bootstrapping. Specifically, individual trials were randomly resampled with replacement to generate a resampled trial sequence, which was refitted using the qCSF procedure. We repeated this procedure of resampling and refitting 10,000 times to generate bootstrap distributions of the fitted parameters, along with associated confidence intervals. To assess statistical significance for differences in qCSF parameters and SF sensitivity estimates between groups, we computed the difference from the 10,000 random pairs of values from the bootstrap distributions of the two groups, and defined p-values as the proportion of samples that "crossed" zero. Note that for all of the key comparisons in the results significant p-values also correspond to non-overlapping confidence intervals.

***Equivalent noise paradigm.*** We used an equivalent noise paradigm (21-25) where perceptual thresholds are measured as a function of varying external noise levels added to the stimuli. Contrast thresholds were measured using a 2-AFC orientation discrimination task, for various amount of external noise (**Fig.4a**). Each trial began with a dynamic fixation point. After a blank screen, a ±45º oriented Gabor signal (cosine envelope diameter: 2 dva) embedded in different intensity levels of dynamic white noise (8 levels, from 0 to .33 SD) was presented at fovea for 100 ms. Participants were asked to report the orientation (±45º from the vertical) of the Gabor patch on each trial. Each stimulus was accompanied with a brief tone to reduce temporal uncertainty and auditory feedback was provided for both correct and incorrect responses. Stimulus presentation was controlled using the *FAST* method (29), an advanced adaptive psychophysical technique used to accurately estimate relevant model parameters in just 480 trials per participant for each SF. This allowed us to estimate contrast thresholds as a function of external noise contrast levels for both 70.71%- and 79.37%-correct difficulty levels, similar to previous studies (21,22,24,25). Five different SFs (0.5, 1, 3, 9 and 16 cpd) were tested across different experimental sessions of 480 trials each (divided into 4 blocks of 120 trials). Data from the equivalent noise experiment were pooled from the *FAST* structures to estimate psychophysical thresholds for each participant at each of the external noise levels (60 trials per level). This approach yielded independent, albeit noisy, contrast threshold estimates at each noise level. Thresholds were estimated by fitting a Weibull function at each external noise level independently:

$$P(c) = 1 - (1 - 0.5) * 2^{-\left(\frac{\log(c)}{\alpha}\right)^{\eta}} \qquad (3)$$

where $P$ denotes percent correct, $c$ is stimulus contrast, $\alpha$ is contrast threshold at 75%-correct performance level, and $\eta$ is the slope of the function. Similar to Park and colleagues (24), we used a Bayesian model fitting method implementing a Markov Chain Monte Carlo (MCMC) technique to estimate the two free parameters $\eta$ and $\alpha$. Specifically, we sampled the posterior distributions of the parameters using JAGS sofware (http://mcmc-jags.sourceforge.net). We assumed a broad uniform prior on each parameter with a range that includes all practically possible values. Maximum a posteriori (the mode of the posterior) were used as the best estimates of the model parameters. We discarded the first 15,000 samples as a burn-in period, and thinned the samples to reduce correlations by only selecting every 200 samples. A total of 10 chains were run in parallel, resulting in 1,000 posterior samples per chain. Thresholds at 70.71% and 79.37% were then computed from the estimated Weibull functions.

**Conventional Perceptual Template Model (PTM) analysis.** The PTM model (21,22) was used to estimate the sources of signal-to-noise changes responsible for performance differences between KC and typical observers. Contrast thresholds for two difficulty levels (70.71% and 79.37%) and across external noise levels were fitted with the PTM to quantify the effects of noise on contrast perception. The PTM consists of five main components (**Fig.4b**): a perceptual template tuned to the signal, a non-linear transducer function, a multiplicative internal noise source ($N_{mul}$), an additive internal noise source ($N_{add}$), and a decision process. The output of the perceptual template is processed by two pathways: the signal pathway in which the output is processed by an expansive nonlinear transducer function, and the multiplicative internal noise pathway in which the output is processed by a rectified nonlinear transducer function. Multiplicative noise is an independent noise source whose amplitude is proportional to the (average) amplitude of the output from the perceptual template, acting as a contrast gain control mechanism. Additive internal noise is another noise source whose amplitude does not vary with signal strength, and is related to the gain of the perceptual template. Both multiplicative and additive noises are added to the output from template matching, and the noisy signal is submitted to a decision process. Here, contrast thresholds ($C_{\mathcal{T}}$) are characterized by:

$$C_{\mathcal{T}} = \frac{1}{\beta}\left[\frac{(1+N_{mul}^2)\,N_{ext}^{2\gamma} + N_{add}^2}{(1/d'^2 - N_{mul}^2)}\right]^{\frac{1}{2\gamma}} \qquad (4)$$

where the input (signal + external noise $N_{ext}$) is filtered through a perceptual template, resulting in signal enhancement via a gain factor $\beta$. The output of the filter is then transformed through a nonlinear transducer function that amplifies the inputs to the $\gamma^{th}$ power, with both internal additive noise ($N_{add}$) and internal multiplicative noise ($N_{mul}$) being added to the output. $N_{add}$ remains constant across signal levels, while $N_{mul}$ is proportional to the signal strength. $N_{ext}$ is manipulated by the experimenter along with the input signal. Finally, a decision process determines the contrast threshold at a specific performance level ($d'$).

The PTM (21,22) considers that performance differences result from changes in three possible sources of inefficiency (**Fig.4b,c**): 1) changes in internal additive noise ($A_{add}$), reflecting signal enhancement of the perceptual template; 2) changes in external noise filtering ($A_{ext}$), corresponding to changes in signal selectivity (tuning) of the perceptual template; and 3) changes in internal multiplicative noise ($A_{mul}$), acting as a contrast gain control mechanism compressing the perceptual template's responses to signal contrast. Group averages of the independently estimated thresholds were used to fit the conventional PTM across SFs. To characterize group differences between NT and KC observers at each SF, the PTM introduces three coefficient indices ($A_{mul}$(SF,group), $A_{add}$(SF,group), and $A_{ext}$(SF,group)) to Equation (4):

$$C_{\mathcal{T}} = \frac{1}{\beta}\left[\frac{\left(1+\left(A_m(SF,group)*N_{mul}(SF)\right)^2\right)\left(A_e(SF,group)*N_{ext}\right)^{2\gamma}+\left(A_a(SF,group)*N_{add}(SF)\right)^2}{\left(\frac{1}{d'^2}-\left(A_m(SF,group)*N_{mul}(SF)\right)^2\right)}\right]^{\frac{1}{2\gamma}} \qquad (5)$$

The coefficient indices for NT are fixed to 1 ($A_{mul}$(NT) = $A_{add}$(NT) = $A_{ext}$(NT) = 1). $N_{add}$(SF) and $N_{mul}$(SF) did not vary across groups, only across SFs to reflect differences in contrast thresholds related to stimulus SF. To estimate whether and how these three types of noise could account for group differences in KC group(s), we computed the relative differences in the effects of each source of noise between the groups. Elevated sources of noise in KC relative to NT would correspond to $A_{mul}$(KC), $A_{add}$(KC), and/or $A_{ext}$(KC) estimates being higher than 1 (i.e., NT group), and to poorer contrast thresholds. Conversely, reduced sources of noise in KC relative to NT would correspond to estimates lower than 1, and to better contrast thresholds. These three coefficient indices for each KC group could be restricted, either varying or fixed at 1. We compared the results obtained by fitting the eight possible models (**Fig.S5** and **Table S2**), ranging from no change in any of the three noise-reduction mechanisms in KC groups to the full model with changes in all three noise-reduction mechanisms. We compared the NT group to 2 groups of KC participants (mild/moderate and severe) at 5 different SFs. The full model consisted of 12 shared parameters across groups (β, γ, $N_{mul}$(SF1-5), $N_{add}$(SF1-5)), and fifteen noise parameters ($A_{mul}$(SF1-5) = $A_{add}$(SF1-5) = $A_{ext}$(SF1-5)) for each of the two KC groups, for a total of 42 parameters. Model fitting was carried out using a least-square method, and goodness-of-fit ($r^2$) for each candidate model was computed as follows:

$$r^2 = 1 - \frac{\sum\left[\log\left(C_{\mathcal{T}}^{predicted}\right) - \log\left(C_{\mathcal{T}}\right)\right]^2}{\sum\left[\log(C_{\mathcal{T}}) - mean(\log(C_{\mathcal{T}}))\right]^2} \qquad (6)$$

where $\sum$ and $mean$ were computed across groups, SFs, external noise levels, and difficulty levels. Given the large differences in the number of free parameters across models, we computed the Bayesian Information Criterion (BIC) as criterion for model selection. The BIC selects the best model while correcting for overfitting by introducing a penalty term for the number of free parameters in the model:

$$BIC = \ n * \log\left(\frac{RSS}{n}\right) + k * \log(n) \qquad (7)$$

where $n$ being the number of predicted data points, $RSS$ is the residual sum of squares of the model, and $k$ is the number of parameters of the model. BIC approximates a transformation of a model's posterior probability. We then computed ΔBIC as:

$$\Delta BIC^k = \ BIC^k - BIC^{k^*} \qquad (8)$$

where $k^*$ correspond to the model with the lowest BIC value (i.e., the best model).

**Hierarchical Bayesian PTM analysis.** To better assess for links between habitual optical quality and individual variability in the PTM noise estimates, we used a hierarchical Bayesian modeling technique to fit the PTM to each participant. This technique assumes that each participant is drawn from a population distribution (NT or KC), which increases statistical power. Importantly, this technique allows estimation of PTM parameters for each participant within a population, providing a better understanding of individual variability. We assumed that individual's response for a given trial is drawn from a Bernoulli distribution (i.e., $response_{ijk} \sim \ Bernoulli(\theta_{ijk})$), where $i$ is the difficulty level, $j$ is the external noise level ($N_{ext}$), and $k$ is the trial number. $\theta_{ijk}$ corresponds to the probability of making a correct response:

$$\theta_{ijk} = 1 - (1 - 0.5) * e^{\left(\frac{m_i \log(x_{ijk})}{\log(C_{\tau ij})}\right)^{\eta}} \qquad (9)$$

where $x$ corresponds to the stimulus contrast, $\eta$ to the slope, $C_{\tau ij}$ to the contrast threshold, and $m_i$ to:

$$m_i = -\log\left(\frac{1-q_i}{1-0.5}\right)^{\frac{1}{\eta}} \qquad (10)$$

The parameter $q_i$ corresponds to the difficulty level (i.e., either .7071 or .7937). The PTM defined the contrast threshold $C_{\tau ij}$ as:

$$C_{\tau ij} = \frac{1}{\beta}\left[\frac{(1+N_{mul}^2)\left(w_{ext}\, N_{ext_{ij}}\right)^{2\gamma} + N_{add}^2}{(1/d'^2_i - N_{mul}^2)}\right]^{\frac{1}{2\gamma}} \qquad (11)$$

Equation (11) is the same as Equation (4), except for the fact that a coefficient for external noise ($w_{ext}$) is added here to characterize the impact of external noise on individual's contrast thresholds. For each participant, we estimated three parameters $N_{mul}$, $N_{add}$, and $w_{ext}$. We assumed a fixed β (1.25) and γ (2) for all participants to simplify the model, which was within a reasonable range reported in previous studies (21,22,24). Similar to the Weibull fitting procedure, an MCMC technique was used to sample from the posterior distributions and estimate the free parameters. Here, we assumed hierarchical priors on each of the model parameters, meaning that each model parameter that characterized an individual participant was assumed to be drawn from an independent Gaussian population distribution with a mean and SD that characterized each group (NT or KC). Priors for population means and SDs were set to broad uniform distributions. Maximum a posteriori were used as the best parameter estimates for both individual and population parameters. We used 15,000 iterations for burn-in, and only selected every 200 samples for thinning. Ten chains were run in parallel, each of which sampled 2,000 posterior samples.

### SI Results

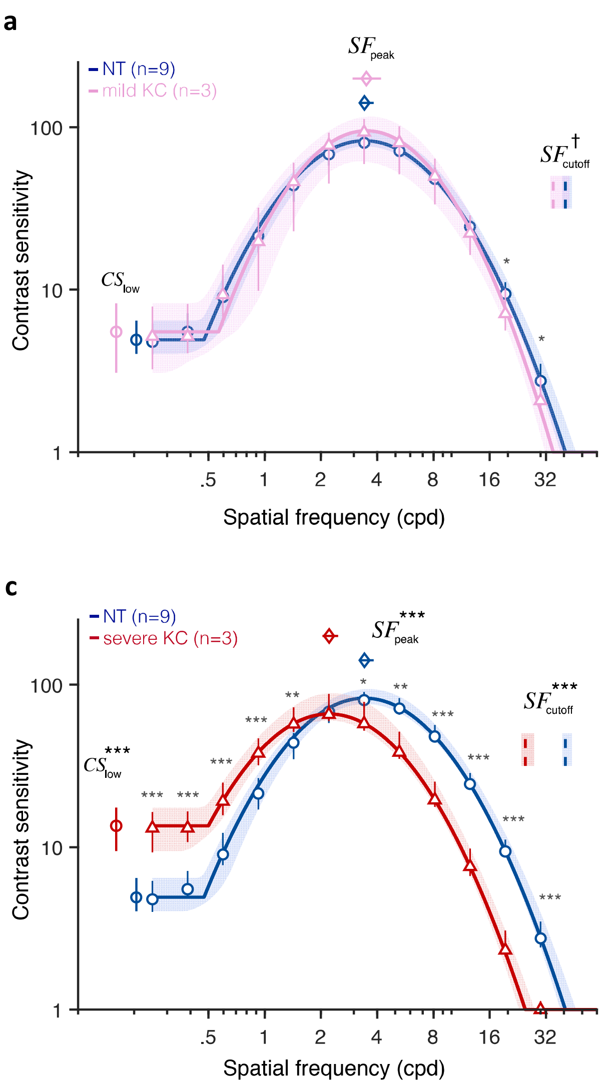


**Figure S3.** *Altered contrast sensitivity function (CSF) following long-term neural adaptation to poor optics.* qCSF results under full AO correction for neurotypical participants (NT; blue curve) and KC participants with either **(a)** mild (N=3), **(b)** moderate (N=3), or **(c)** severe (N=3) habitual optical aberrations. Relative to NT participants, KC observers showed altered CSF functions that were shifted towards lower SFs, with impaired sensitivity at high SFs as well as improved sensitivity at low SF. This pattern depended on KC severity, with mild KC showing almost no difference compared to NT observers while patients with severe KC showed marked CSF alterations. Shaded areas and error bars represent bootstrapped 95%-CI. Asterisks indicate significant differences computed from bootstrapping between NT and KC participants ($^{\dagger}$: *p<0.1;* *: *p<0.05*; **: *p<0.01*; ***: *p<0.001*).

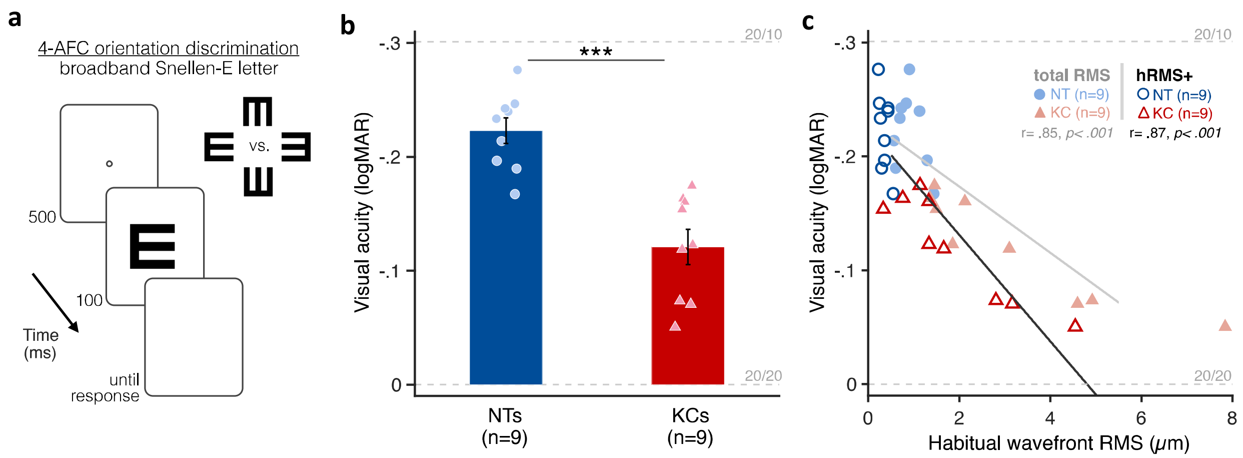


**Figure S4.** *Poorer high contrast visual acuity (VA) following long-term neural adaptation to poor optics.* **(a)** Visual acuity was measured under AO correction using a 4-alternative forced choice discrimination task. The size of Snellen E letter stimuli varied from trial-to-trial to estimate 62.5%-correct VA thresholds in logMAR. **(b)** Relative to neurotypical (NT) observers, KC patients showed poorer high contrast VA despite being tested under similar aberration-free optical condition. Error bars correspond to ±1 SEM with each data point representing an individual observer. **(c)** Poorer habitual optical quality (total RMS or hRMS+) was associated with impaired VA when tested under aberration-free conditions.

### Experiment 2: Model comparisons

**Table.S2** summarizes the results from each of the 8 variants of the PTM, with **Fig.S5** showing the fits from each model. To account for large differences in the number of free parameters across models, we computed the Bayesian Information criterion (BIC) to evaluate which model best fits the data while penalizing for greater number of free parameters. We subtracted the lowest BIC value corresponding to the best model to compute the ΔBIC (**Table S2** and **Fig.5b**). The internal additive noise ($A_{add}$) model was the best model (**Fig.S5b**), followed by mixture models that included internal additive noise as one of the free parameter (**Fig.S5c,d**), and by the full model (**Fig.S5a**). Other models that did not allow changes in internal additive noise ($A_{add}$) were particularly poor and could not account for the differences between groups (**Fig.S5e-h**). These results support the fact that SF-specific changes in internal additive noise mediates the gains and losses in sensitivity across SFs in KC patients with severe optical defects.

**Table S2.** Summary table of the different variants of the Perceptual Template Model (PTM)

| Model | number of parameters (k) | variance explained ($r^2$) | residual sum of squares (rss) | Δ BIC |
|---|---|---|---|---|
| null | 12 | 82.52% | 5.59 | 196.26 |
| $A_{add}$ | 22 | 93.86% | 1.96 | 0 |
| $A_{ext}$ | 22 | 85.70% | 4.57 | 202.94 |
| $A_{mul}$ | 22 | 91.87% | 2.60 | 67.46 |
| $A_{add}$ + $A_{ext}$ | 32 | 94.96% | 1.61 | 7.32 |
| $A_{add}$ + $A_{mul}$ | 32 | 94.92% | 1.62 | 9.11 |
| $A_{mul}$ + $A_{ext}$ | 32 | 93.48% | 2.08 | 69.20 |
| full | 42 | 95.70% | 1.37 | 24.19 |

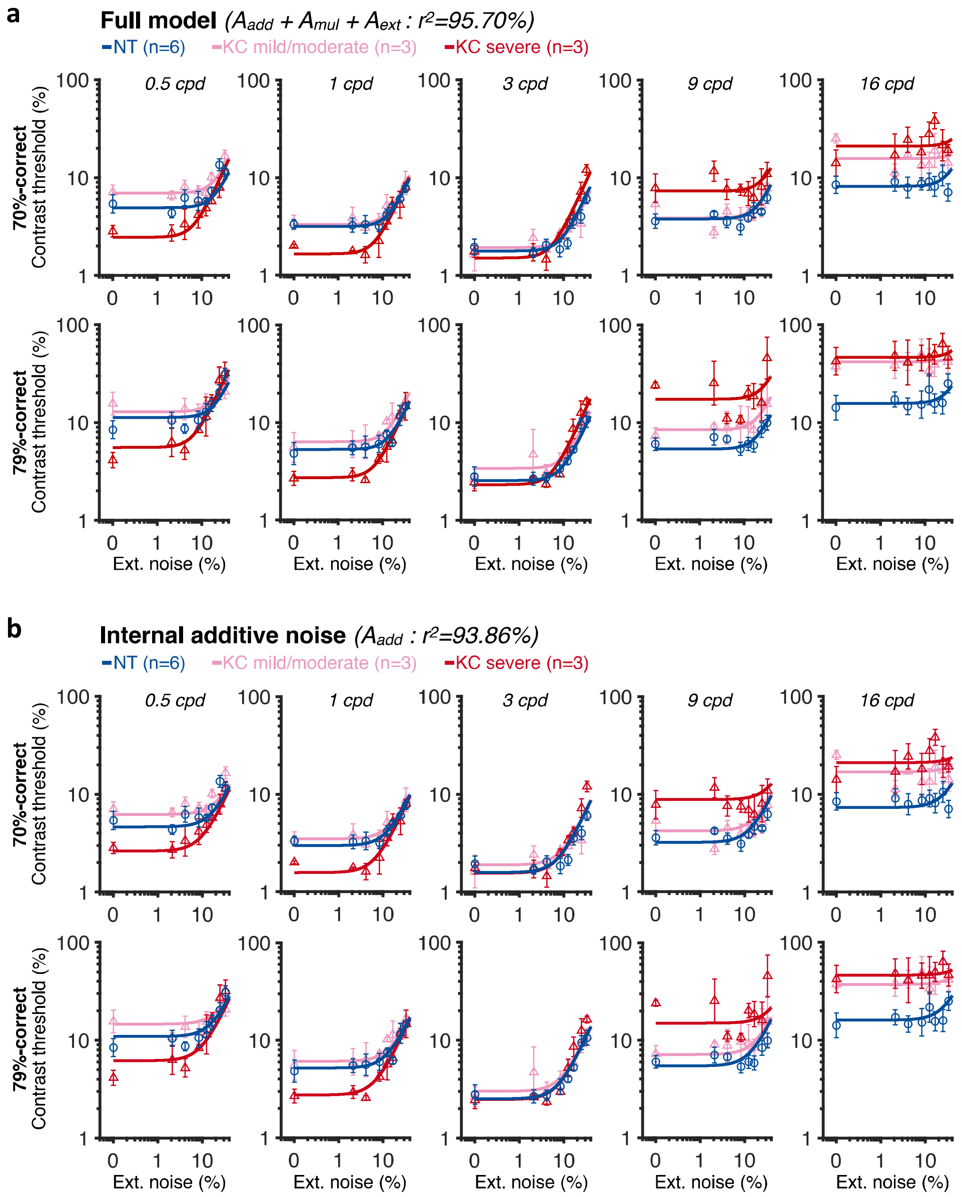


**Figure S5.** *Experiment 2: each variant of the PTM (1/4).* Contrast threshold estimates were measured under AO correction for Gabor stimuli embedded in different levels of external white noise. Threshold-vs-noise functions were obtained for five different spatial frequencies (0.5, 1, 3, 9, and 16 cpd), and at either 70.71% and 79.37%-correct difficulty levels. The PTM was used to evaluate the contribution of distinct sources of inefficiencies. Results for **(a)** the full model (internal additive noise + internal multiplicative noise + external noise filtering; $A_{add}$ + $A_{mul}$ + $A_{ext}$); **(b)** the internal noise model ($A_{add}$); **(c)** the internal additive noise and external noise filtering model ($A_{add}$ + $A_{ext}$); **(d)** the internal additive and multiplicative noise model ($A_{add}$ + $A_{mul}$); **(e)** the internal multiplicative noise and external noise filtering model ($A_{add}$ + $A_{ext}$); **(f)** the internal multiplicative noise model ($A_{mul}$); **(g)** the external noise filtering model ($A_{ext}$); and **(h)** the null model. Each symbol represents average contrast thresholds for each group ±1 SEM error bars.

**c** **Internal additive noise + external noise filtering** *($A_{add} + A_{ext}$ : $r^2$=94.96%)*

-NT (n=6) -KC mild/moderate (n=3) -KC severe (n=3)

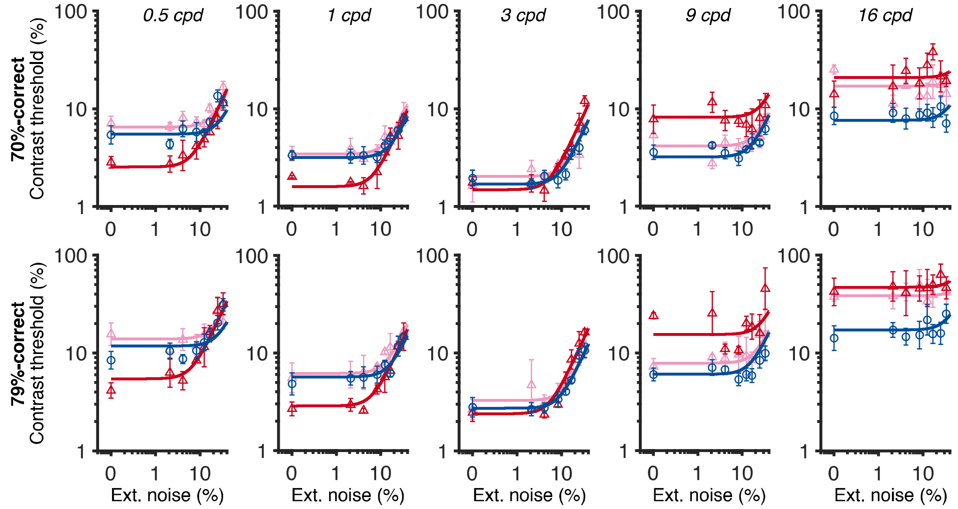


**d** **Internal additive noise + internal multiplicative noise** *($A_{add} + A_{mul}$ : $r^2$=94.92%)*

-NT (n=6) -KC mild/moderate (n=3) -KC severe (n=3)

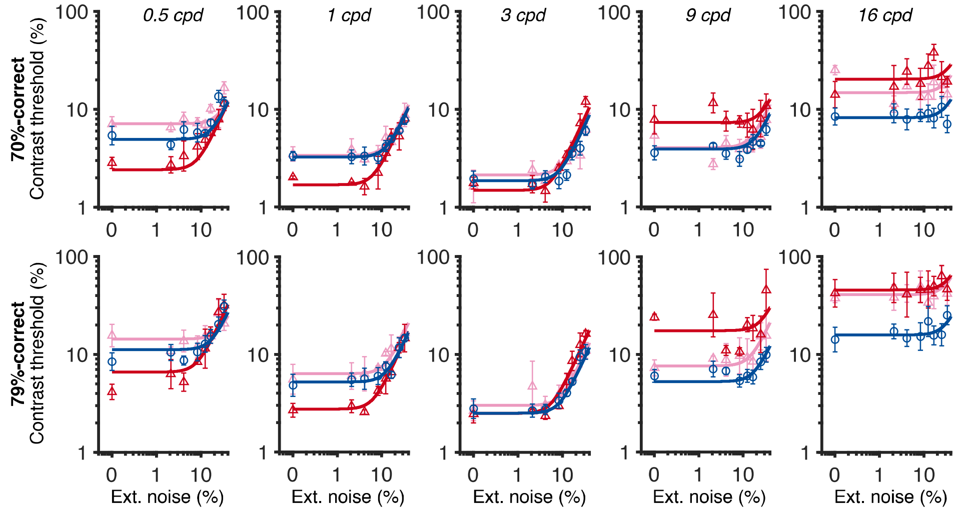


**Figure S5** *(continued)*. *Experiment 2: each variant of the PTM (2/4)*

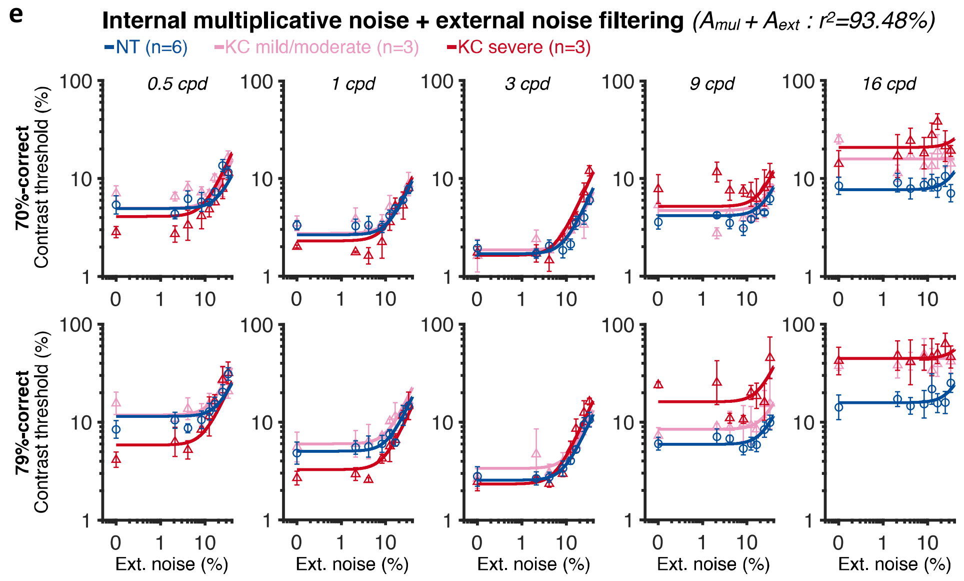


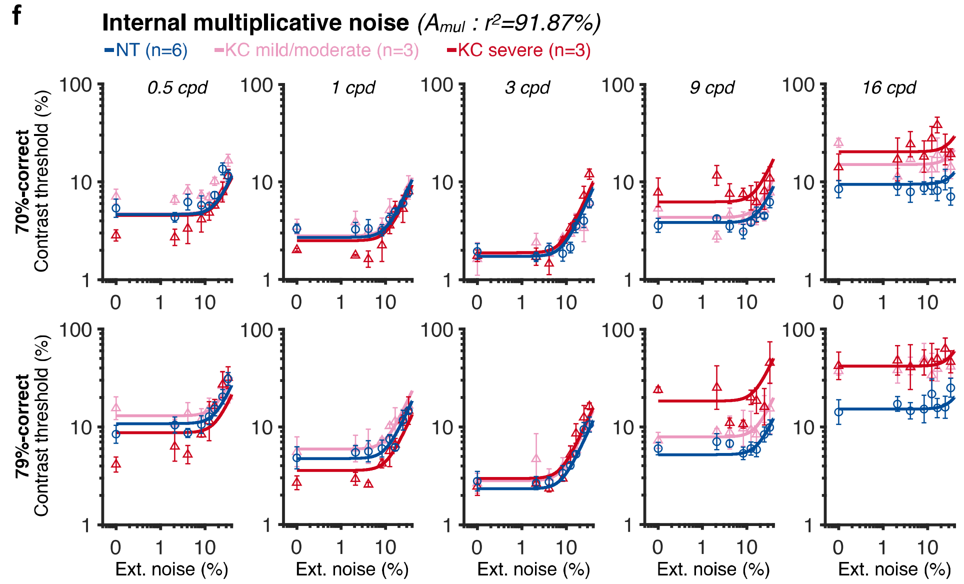


**Figure S5** *(continued)*. *Experiment 2: each variant of the PTM (3/4)*

**g**

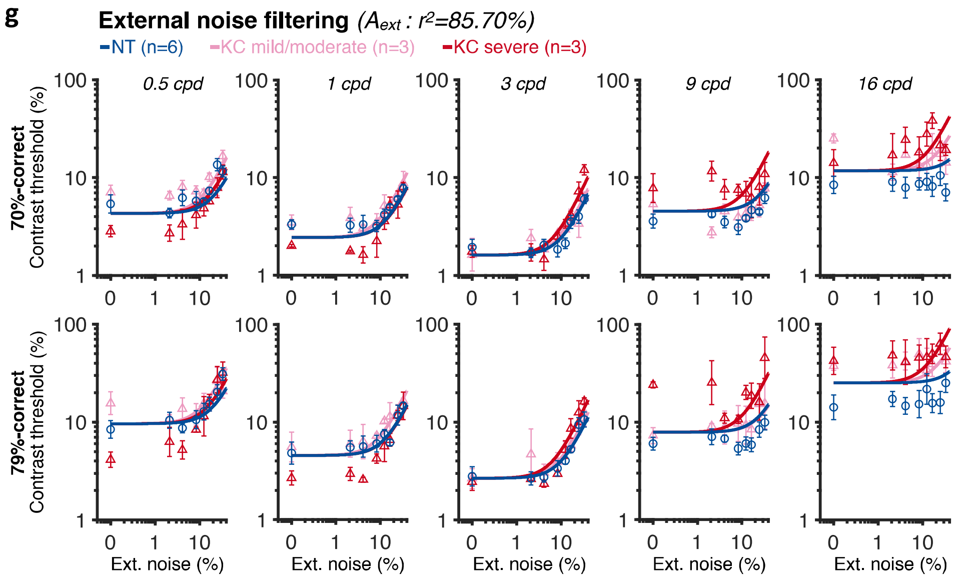


**h**

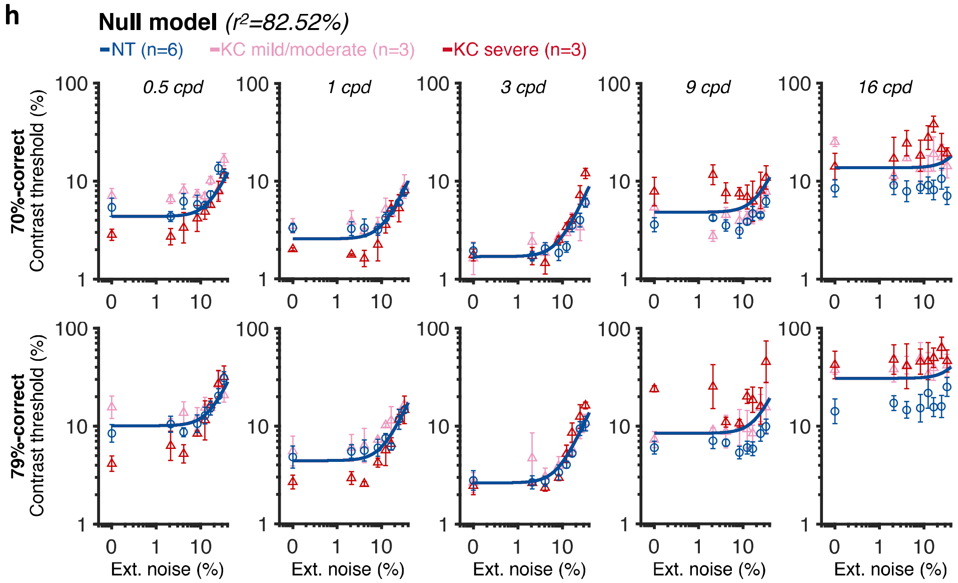


**Figure S5** *(continued)*. *Experiment 2: each variant of the PTM (4/4)*

### Experiment 2: Independent PTM analyses for low and high spatial frequency conditions

Conventional PTM analyses across all five SFs revealed that changes in internal additive noise ($A_{add}$) could account for the differences in contrast thresholds between NT observers and KC patients (**Fig.5**). To further assess the mechanisms underlying the pattern of gains and losses in contrast processing observed in KC patients, we ran separate PTM analyses for low SFs (**Fig.S6a,b**; 0.5 and 1 cpd), and for high SFs (**Fig.S6c,d**; 9 and 16 cpd). In both cases, we found that changes in internal additive noise (i.e., signal enhancement) was the model that best explained the SF-specific differences in contrast thresholds across external noise levels and difficulty levels:

-at low SFs (**Fig.S6a**; 0.5 and 1 cpd), changes in internal additive noise explained most of the differences across groups, with the internal additive noise model ($r^2$=94.73%) was significantly better than the null model ($r^2$=81.96%; $F_{4,86}$=57.35, *p<0.001*). Although it was different from the full model ($r^2$=95.81%; $F_{8,78}$=3.07, *p=0.005*) due to the higher number of free parameters, the Bayesian Information Criterion indicated that the internal additive noise model was the best model explaining the differences in contrast thresholds between groups (**Fig.S6b**). Relative to NT participants, severe KC participants showed reduced internal additive noise ($A_{add}$: -41.33% and –43.92% at 0.5 and 1 cpd, respectively), whereas moderate KC showed elevated internal additive noise ($A_{add}$: +31.77% and +15.44% at 0.5 and 1 cpd, respectively) relative to our NT group.

- at high SFs (**Fig.S6c**; 9 and 16 cpd), the best model remained a model assuming solely differences in internal additive noise. This model ($r^2$=92.23%) was significantly better than the null model ($r^2$=68.76%; $F_{8,82}$=71.44, *p<0.001*). Although it was different from the full model ($r^2$=93.79%; $F_{4,78}$=2.99, *p=0.006*), model comparisons indicated that the internal additive noise model was the best model (**Fig.S6d**). Relative to NT participants, KC participants showed increased internal additive noise by +32.86% and +125.17% at 9 and 16 cpd, respectively for mild/moderate KC, and by +172.86% and +179.13% for severe KC at 9 and 16 cpd, respectively. Note that contrast threshold estimates were particularly variable and poor at high SFs, which may have limited our ability to observe potential differences in external noise filtering and internal multiplicative noise in addition to the primary elevation in internal additive noise observed at high SFs.

In summary, our results strongly support changes in internal additive noise as the primary mechanism underlying differences in contrast sensitivity at both low SFs and high SFs.

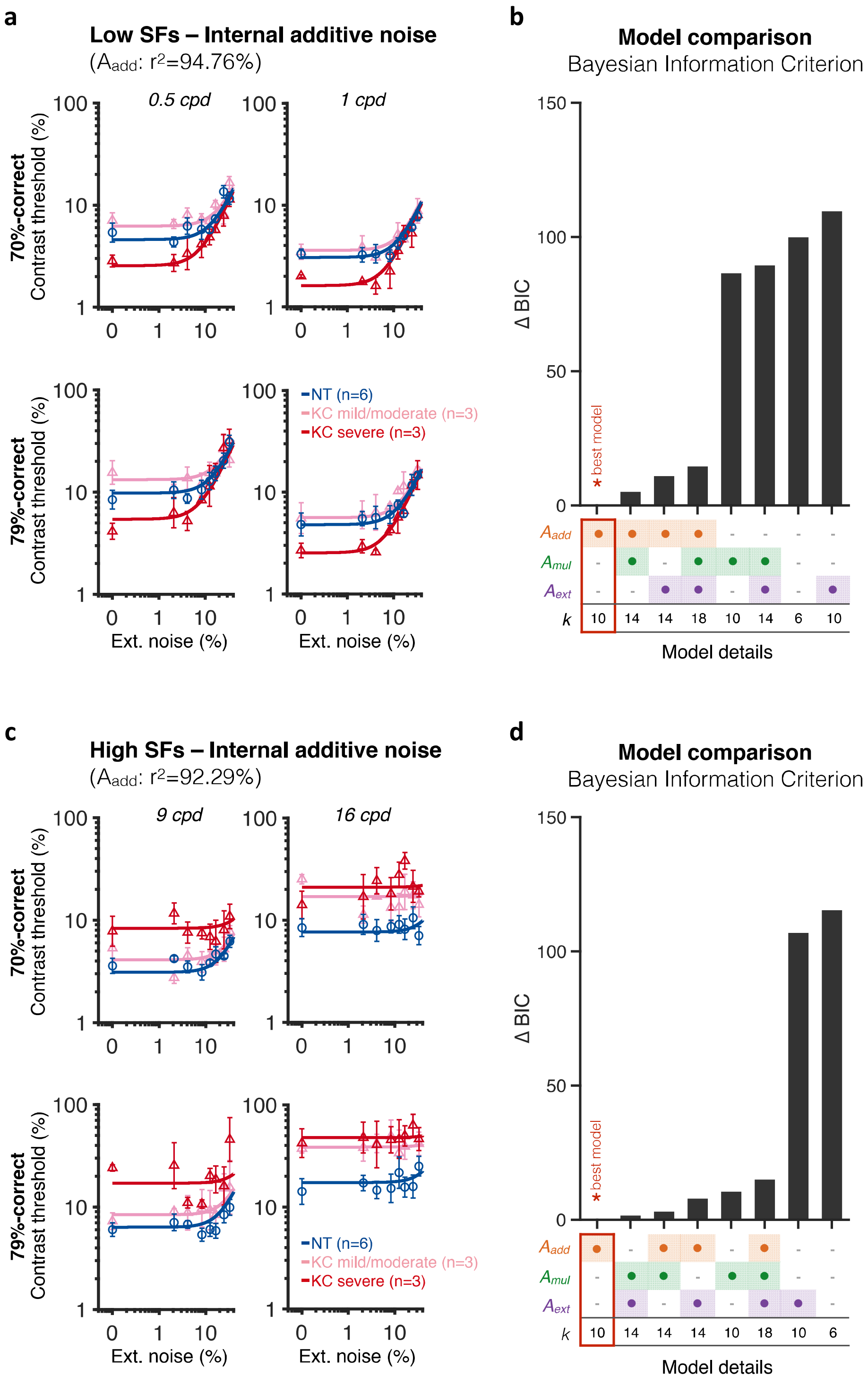


**Figure S6.** *Independent PTM analyses for low and high spatial frequencies (SF) support the internal additive noise model.* At both **(a)** low SFs and **(c)** high SFs, differences in contrast thresholds between NT and KC observers were best explained by differences only in internal additive noise ($A_{add}$; low SFs: $r^2$=94.76%; high SFs: $r^2$=92.29%), reflecting improved signal enhancement in patients with severe KC. **(b,d)** Bayesian Information Criterion (BIC) evidence was computed for each model and the value from the best model was subtracted. The best model (ΔBIC = 0) for both low and high SFs was a model assuming changes only in internal additive noise between NT and KC observers, consistent with the results across SFs.

### Experiment 2: Contrast sensitivity at low and high external noise levels and contrast threshold ratios between difficulty levels

**Fig.S7** shows contrast sensitivity (1/contrast threshold) computed either for the lowest two external noise levels (**Fig.S7a**), or for the two highest external noise levels (**Fig.S7b**) tested. Consistent with the pattern of gains and losses in contrast sensitivity observed in the qCSF experiment (**Fig.3** and **Fig.S3**), observers with severe KC showed enhanced sensitivity for low SFs and impaired sensitivity for high SFs when almost no external noise was added to the target stimuli. This pattern was less pronounced in mild/moderate KC participants. Note that this result indicates that the alterations in qCSF observed in Experiment 1 were not due to the larger uncertainty regarding the SF of the upcoming target stimulus on a given trial, as a similar pattern was observed in Experiment 2 for which the SF of the signal did not change during an entire experimental sessions. At high external noise levels, contrast sensitivity was more comparable across groups, which is consistent with the predictions of the internal additive noise model (**Fig.4c**). Differences at high SFs under high external noise are likely due to the fact that the nonlinearity characteristic of TvN curves was not well captured and internal additive noise most likely remained the main limiting factor even at the high external noise levels.

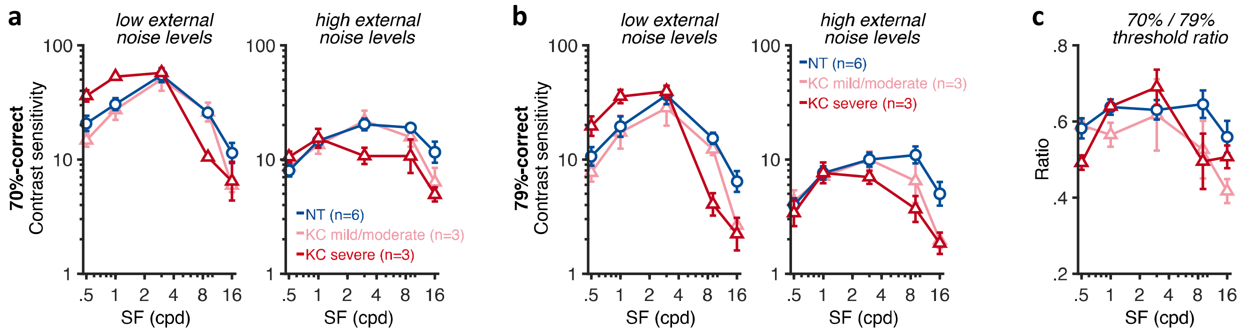


**Figure S7.** *Contrast sensitivity at low and high external noise levels and contrast threshold ratios between difficulty levels.* **(a,b)** Contrast sensitivity (1/threshold) was computed as a function of the stimulus spatial frequency (SF) using either the two lowest external noise values (*left panels*) or the two highest external noise values (*right panels*), at **(a)** 70.71%-correct and **(b)** 79.37%-correct levels of difficulty. Each symbol represents average values for each of the three groups, with ±1 SEM error bars. A similar pattern was observed at the two difficulty levels. At low external noise, SF-specific differences in contrast sensitivity were observed, consistent with the alterations in qCSF observed in Experiment 1. **(c)** Contrast threshold ratios between difficulty levels were computed to assess a possible role of internal multiplicative noise. Although it seemed that larger differences were observed with difficulty levels between NT and KC participants at high SFs, this pattern was likely due to the particularly high variability in threshold estimates observed at high SFs. Additional analyses ruled out differences in internal multiplicative noise as a factor mediating the changes in contrast sensitivity observed between NT and KC participants.

To test for the possible influence of internal multiplicative noise, we computed the ratio of contrast thresholds at the two difficulty levels (i.e., 70.71% and 79.37% correct), averaged across external levels (**Fig.S7c**). Similar ratios between groups would indicate comparable differences in contrast threshold across difficulty levels between NT and KC participants, which would rule out differences in internal multiplicative noise ($A_{mul}$; **Fig.4c**). An ANCOVA on the ratios of thresholds across external noise with participants' habitual (total) RMS as a covariate indicated a marginal main effect of SF ($F_{4,40}$=2.52, *p=0.056*, $\eta_p^2$=.20), with a significant interaction between SF and participant's habitual RMS

($F_{4,40}$=3.60, *p=0.013*, $\eta_p^2$ =.27). The same trend was observed using hRMS+ as a measure of the participant's habitual optical quality. Similarly, when comparing ratios between NT and KC groups, we found a marginal interaction between SF and group ($F_{8,36}$=2.06, *p=0.067*, $\eta_p^2$ =.31), suggesting that KC groups showed larger differences in thresholds between difficulty levels at high SFs. Several factors must be taken into account when considering internal multiplicative noise as a potential subsidiary source of inefficiencies. First, the internal additive noise model was the best model explaining the differences in contrast thresholds between groups, regardless of whether we analyzed all SFs together (**Fig.5**) or separately analyzed low SFs and high SFs (**Fig.S6**). Second, individual difference in internal additive noise was the only parameter correlating with the magnitude of habitual optical aberrations of each participant (**Fig.S8**), which supports the finding that SF-specific changes in signal enhancement mechanisms alone can account for the impact of long-term exposure to poor optical quality on visual processing. The PTM has been successfully used to characterize the mechanisms underlying differences in visual functions for a wide range of brain functions, such as attention (30, 31) and perceptual learning (21, 22), as well as between specific populations, such as in amblyopia (32), autism (24), and dyslexia (33). It is possible that the apparent differences in ratios across difficulty levels at high SFs may be due to the fact that contrast thresholds were particularly elevated and variable in KC patients for high SFs. Neural insensitivity at high SFs also limited our ability to capture the nonlinear aspect characteristic of TvN curves, which could have affected the ability of the PTM to identify differences in external noise filtering and/or internal multiplicative noise as potential secondary source of inefficiency. In summary, our results clearly indicate that the effects of long-term exposure to optical defects on contrast sensitivity at different SFs are mediated primarily by changes in internal additive noise.

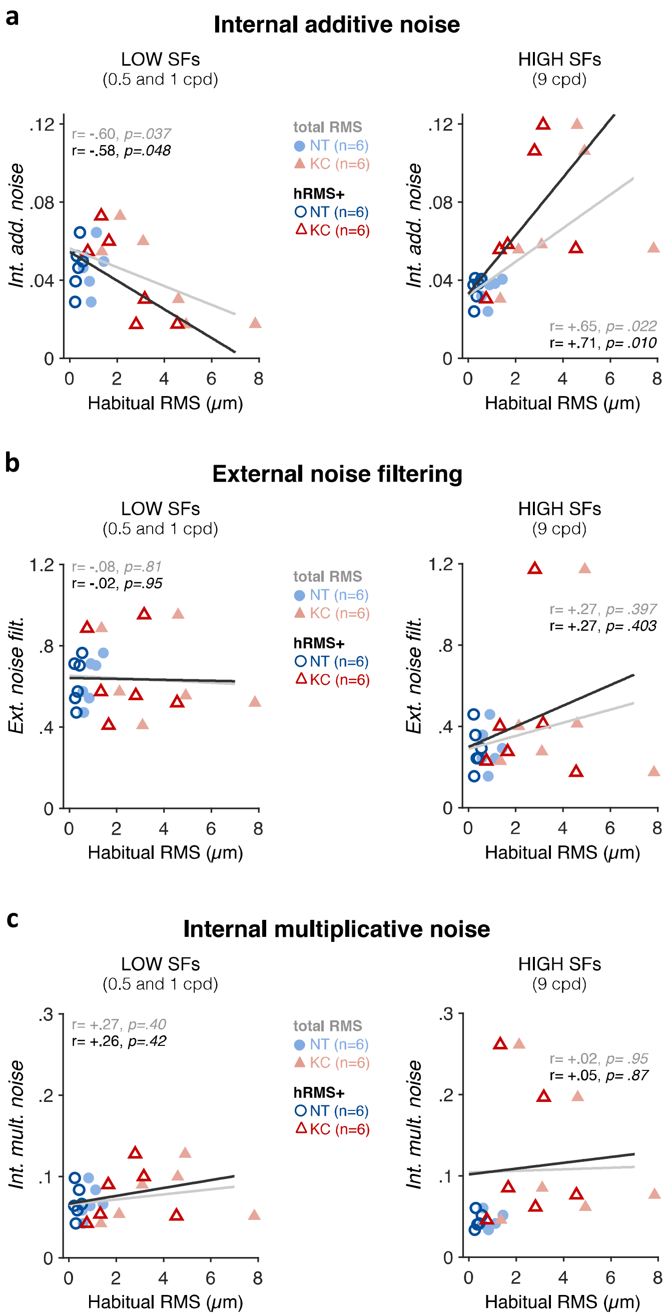


**Figure S8.** *Correlation between individual PTM estimates and habitual optical quality at low and high SFs.* **(a)** Internal additive noise ($A_{add}$). Poorer habitual optical quality was associated with reduced internal additive noise at low SF (*left panel*) and elevated internal additive noise at high SF (*right panel*). Consistent with internal additive noise being the primary source of inefficiency mediating the effects of long-term exposure to poor optical quality, habitual optical quality did not correlate with either **(b)** external noise filtering ($A_{ext}$) or **(c)** internal multiplicative noise ($A_{mul}$).
